# GATOR2 regulates the nutrient-dependent recruitment of GATOR1 to lysosomes

**DOI:** 10.64898/2026.07.07.737070

**Authors:** Yingbiao Zhang, Chun-Yuan Ting, Shu Yang, Richard Garcia, Mary Lilly

## Abstract

TORC1 is a central regulator of cellular metabolism. TORC1 activation depends on its recruitment to the lysosome, a process mediated by the Rag GTPases and their upstream regulator, the GATOR complex. GATOR consists of two subcomplexes: GATOR1, which converts the Rag GTPases into an inactive form during nutrient starvation by acting as a GAP towards RagA, and GATOR2, which counteracts GATOR1 through an unknown mechanism. Here we dissect the GATOR-Rag GTPases network at the cellular and subcellular level in Drosophila using genomically tagged proteins. We find that GATOR2 maintains GATOR1 in the GAP-inactive state under nutrient-replete conditions. Moreover, using fluorescent recovery after photobleaching we show that while GATOR1 and GATOR2 are recruited to lysosomes as a supercomplex in fed conditions, their recruitment becomes decoupled during starvation. Taken together, our findings support a model in which GATOR1 forms an inactive supercomplex with GATOR2 under nutrient-rich conditions. However, upon nutrient starvation, this supercomplex dissociates, enabling GATOR1 to adopt a GAP-active state and inhibit the Rag GTPases, thereby preventing the recruitment and activation of TORC1 on lysosomes. Finally, our data identify Wdr59 as a critical regulator of this nutrient-dependent remodeling that is required to relieve GATOR2-mediated inhibition of GATOR1.

## INTRODUCTION

The multiprotein Target of Rapamycin 1 (TORC1) is a key regulator of metabolism in eukaryotic cells ^1-4^. To maintain metabolic balance, TORC1 integrates signals from various pathways that track a broad range of inputs from both inside and outside the cell, such as amino acids, growth factors, ATP, and cholesterol levels ^5^. When growth signals and nutrients are abundant, TORC1 functions as a serine-threonine kinase which activates downstream targets that promote protein synthesis, ribosome production, and other anabolic processes linked to cell growth. In contrast, TORC1 suppresses catabolic processes and autophagy by phosphorylating proteins that drive these activities, including ATG1/ULK1 and members of the MiT/TFE transcription factor family^6–8^. Dysregulation of TORC1 is associated with several human diseases and pathological processes, including cancer, epilepsy, and aging ^9^. Therefore, understanding the pathways that regulate TORC1 is a major area of research.

TORC1 activity is controlled by its intracellular localization ^5^. When growth signals and nutrients are abundant, TORC1 is recruited to lysosomes by the Rag GTPase, where it is activated by the small GTPase Rheb ^10^. The Rag GTPase functions as an obligate heterodimer, consisting of a RagA/B subunit bound to a RagC/D subunit, and is anchored to the lysosomal membrane through the scaffold protein complex, Ragulator ^11–13^. Notably, in *Drosophila*, there is only a single copy of RagA and RagC in the genome ^11^. The activity of the Rag GTPases is regulated by the GATOR complex, originally identified in yeast as the Seh1 Associated (SEA) complex ^14^. Research in both yeast and mammals has shown that the GATOR1 complex, composed of DEPDC5, NPRL2, and NPRL3, acts as GTPase-activating proteins (GAPs) that inhibits TORC1 activity by promoting the conversion of RagA^GTP^ to its inactive RagA^GDP^ state in response to amino acid limitation ^15–18^. In *Drosophila* the DEPDC5 homolog is named Iml1^19^. Whole animal studies in metazoans, including *Drosophila* and mice, further support the essential role of GATOR1 in maintaining metabolic homeostasis and coordinating the response to nutrient stress at both the cellular and organismal levels ^5,19–22^.

The GATOR2 complex opposes the activity of GATOR1, thereby promoting TORC1 activity ^17,19,22,23^. Cryo-electron microscopy (cryo-EM) indicates GATOR2 is a large (1.1 MDa) multiprotein complex consisting of multiple copies of the conserved proteins WDR24, MIOS, WDR59, SEH1L, and SEC13 ^24–26^. Early computational studies in yeast suggested that GATOR2 shares structural features with coatomer proteins and membrane-tethering complexes ^27,28^. This prediction aligns with experimental observations, as GATOR2 is localized to the surface of lysosomes and autolysosomes in *Drosophila* and mammalian tissues, and to the vacuolar membrane in yeast ^17,19,22,23,26,27^. Studies from both yeast and mammals, revealed that the large multiprotein GATOR2 complex, known as SEACAT in yeast, adopts a large two-fold symmetric, cage-like structure ^26,29^. While the overall structure of GATOR2 is conserved from yeast to humans, the mechanism by which GATOR2 opposes GATOR1 activity remains undefined.

Work performed primarily in mammalian tissue culture cells has provided a detailed model for how the GATOR-Rag GTPase-GATOR signaling axis regulates TORC1 in response to amino acid levels ^3,9^. However, it is currently unclear how well this model reflects the regulation of TORC1 activity within the varied developmental and metabolic contexts of a multicellular animal. Notably, in *Drosophila*, the GATOR2 component WDR59 functions as an inhibitor of GATOR2 in most tissues ^22,30^. These findings are consistent with the reported role of Sea3/Wdr59 in fission yeast, as well as observations that deletion of WDR59 in HEK293 cells and mouse embryonic fibroblasts confers partial resistance to nutrient starvation ^31,32^. Taken together these data suggest that the current 1.1 Megadalton cryo-EM structure of GATOR2 and SEACAT may represent the inhibited/inactive form of the GATOR2/SEACAT complex. However, a full understanding of how GATOR2 opposes the activity of GATOR1 to promote TORC1 activation awaits additional functional studies.

While the precise control of TORC1 activity is critical to the growth and development of metazoans, there are limited tools to explore the cell biology of the GATOR-Rag-GTPase-TORC1 signaling axis in vivo. To address this deficiency, we have developed reagents to examine the regulation of the GATOR-Rag GTPase-TORC1 signaling axis in vivo, at the single cell and subcellular level, in the model organism *Drosophila*. The advantage of our system is three-fold (1) It allows for the study of a wide variety of cell types in different developmental and physiological contexts. (2) It uses CRISPR/Cas9 to tag genes at their chromosomal localization, or transgenes that are expressed from their endogenous promoters, to avoid artifacts from over or under expression. (3) It can be used in combination with the powerful molecular genetic tools available in *Drosophila* to examine gene function and complex genetic interactions.

Here we used these newly generated reagents to examine the regulation of the GATOR-Rag GTPase-TORC1 signaling pathway in the *Drosophila* ovary. We demonstrate that many of the hierarchies governing the recruitment of the Rag GTPase and the GATOR complex to lysosomes in mammals are conserved in *Drosophila*. Additionally, we find that in nutrient replete conditions, the GATOR2 complex promotes the retention of GATOR1 in its GAP-inactive, inhibited form. Our lysosomal FRAP data are consistent with GATOR1 and GATOR2 functioning as a supercomplex in nutrient replete conditions, with the lysosomal exchange of the two complexes decoupled in response to nutrient limitation, allowing for the engagement of catalytically active GATOR1 on lysosomes. Notably, FRAP studies support a similar model for GATOR regulation in HeLa cells

## RESULTS

### Reagents to dissect the GATOR-RAG-GTPase signaling pathway in vivo

We are interested in defining the diverse pathways that regulate of TORC1 activity within the complex environment of a multicellular animal. To facilitate tissue specific studies in vivo in the metazoan *Drosophila*, we tagged multiple components of the GATOR-Rag GTPAse-TORC1 signaling pathway at their endogenous loci using CRISPR/Cas9 gene editing with small epitope tags that can be used for localization studies on fixed tissue and biochemical studies, as well as GFP^11^ a GFP fragment that can be reconstituted for live cell imaging (see below) (**Fig. 1A,B**) ^22,33,34^. Importantly, all genomically edited genes were homozygous viable and fertile, including GATOR1 components, *iml1*, *nprl2* and *nprl3* and GATOR2 components *mio*, *wdr24*, and *wdr59*, indicating the associated tagged proteins were functional. Additionally, we generated tagged genomic constructs of the Rag GTPase components RagA and RagC which were also homozygous viable and fertile in that they can rescue *ragA* and *ragC* null mutations respectively ^22^. As predicted, all tagged proteins in the GATOR-Rag GTPAse-TORC1 signaling pathway localized at least in part, to puncta that co-localized with Rab7 and are thus present on lysosomes and autolysosomes (**Fig. 1B-G**).

**Figure 1.**
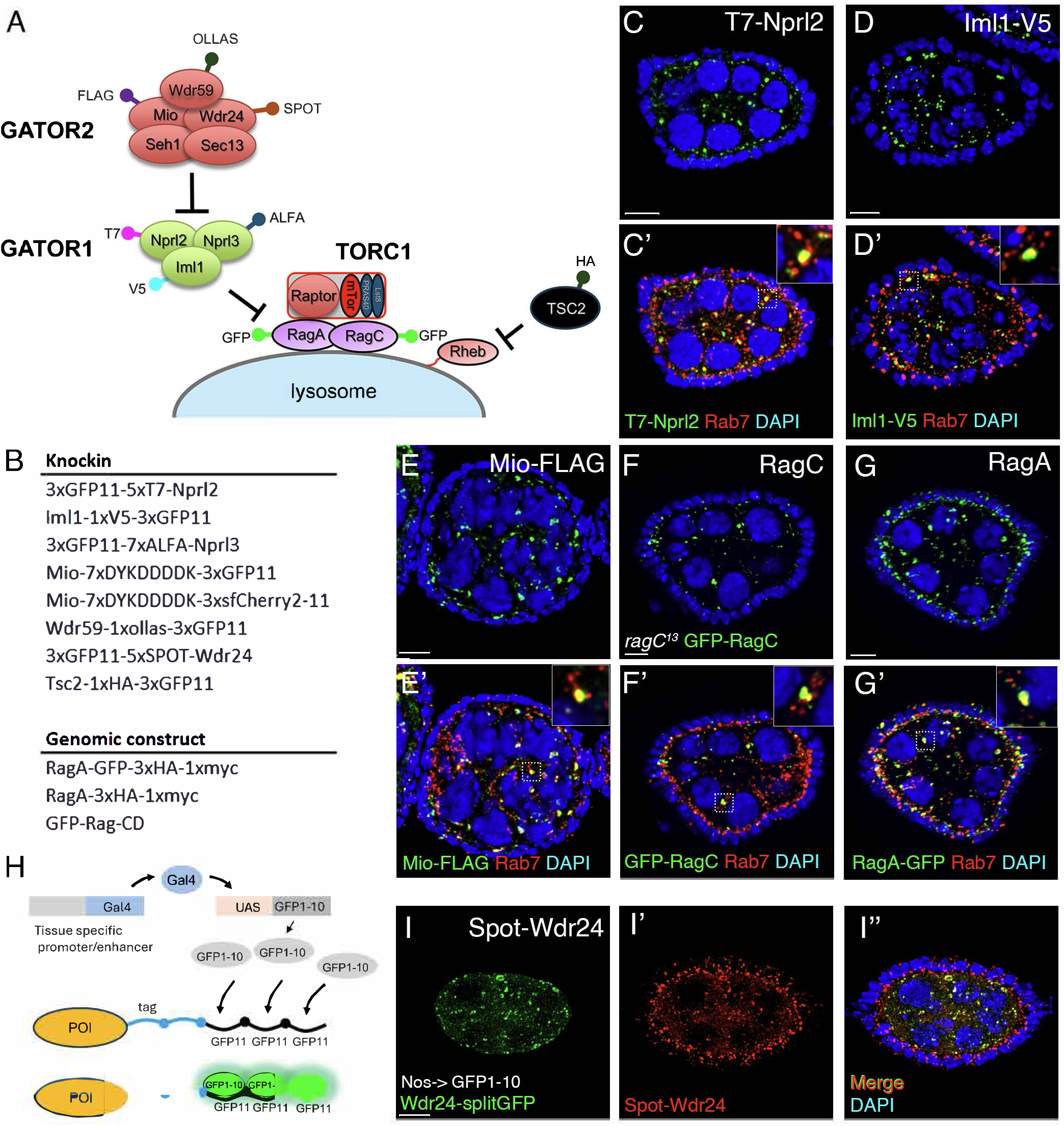
A toolbox for the study of the GATOR-Rag GTPase signaling axis. (A) Schematic illustrates tagged components of the TORC1 signaling pathway. (B) List of knockin lines. (C-G) Immunolocalization of tagged proteins. Ovaries were stained with tag-specific antibodies (green), while the late endosome marker Rab7 (red) was used to label lysosomes. Hoechst 33342 (blue) was used to label DNA. (C, D) GATOR1 subunit, T7-Nprl2 and Iml1-V5. (E) GATOR2 subunit Mio-FLAG. (F) GFP-RagC and (G) RagA-GFP-3HA-1myc. (H) Diagrammatic for split GFP labeling. The protein of interest was tagged with 3xGFP11 and an epitope tag by CRISPR, which only fluoresces upon co-expression of GFP1–10 in the same cell. (I-I’’) Wdr24-rcGFP protein was reconstituted by specifically expressing GFP1-10 using the germline driver, Nos-Gal4. Wdr24-GFP signal was observed only in germline cyst cells, but not in the somatic follicle cells. In contrast, antibodies against the epitope tag Spot, highlights Wdr24-Spot on lysosomes in both germline and somatic cells (I’,I”). Scale bar: 5 μm.

The ability to define the subcellular distribution and behavior of proteins in specific cell types is often compromised by high levels of background staining from surrounding tissues. To avoid this complication, we used the split-GFP tagging strategy ^33^. As noted above, the GFP^11^ tag was added to multiple proteins in the GATOR-Rag GTPase signaling cascade (**Fig 1B**). To visualize the protein of interest in a specific cell type, we expressed split-GFP^1–10^ using UAS-GAL4 tissue specific drivers ^35^. The split-GFP^1–10^ reconstitutes with the GFP^11^ tag on the protein of interest to reveal the subcellular location of the endogenous protein (**Fig. 1H**). As proof of principal, we expressed split-GFP^1–10^ in the female germline using the nosGAL4 UAS driver in females carrying a genomic copy of Wdr24-SPOT-GFP11 ^36^. As anticipated, the lysosomal localization of the Wdr24-GFP11 protein was highlighted by the reconstituted split GFP in the germline but not the somatically derived follicle cells (**Fig. 1l**). In contrast, antibody staining against the SPOT tag reveals the complete distribution of the Wdr24-SPOT-GFP^11^ protein in both germline and somatic cells (**Fig. 1l’)**. In a second example, we visualize endogenously tagged Iml1-GFP^11^ protein in migrating border cells by driving the expression of GFP^1–10^ using the slbo-Gal4 driver **(Supplemental Fig. 1**) ^37^. These data demonstrate that the split-GFP system effectively highlights the subcellular localization of proteins in the GATOR-Rag GTPase signaling pathway in cells where localization might otherwise be obscured by surrounding tissue.

### The Rag GTPase recruits GATOR1 and GATOR2 to lysosomes in *Drosophila*

Recent studies in yeast and mammalian cells support the model that components of GATOR1(SEACIT) and GATOR2 (SEACAT), form a supercomplex that traffics between lysosomes and the cytoplasm ^26,38,39^. In mammalian cells the lysosomal recruitment of GATOR1 and GATOR2 requires both the Rag GTPase, which also traffics to and from the lysosomes, and the multi-protein complex KICSTOR ^38,40^. Notably, the *Drosophila* genome does not encode KICSTOR homologs highlighting potential species-specific differences in this TORC1 regulatory pathway ^40^. Thus, to determine if the regulatory hierarchies that control GATOR-Rag GTPase signaling in mammals is conserved in *Drosophila*, we used in vivo loss of function genetics to determine how individual proteins in the pathway impact the lysosomal recruitment of GATOR2, GATOR1 and the Rag GTPase in *Drosophila*.

To determine if the lysosomal localization of GATOR2 components require a functional Rag GTPase, we generated homozygous clones of a *ragA* null allele, *ragA*^11^, in the germline of females that contained a genomically tagged Mio-FLAG protein. We found that relative to adjacent heterozygous *ragA*^11^/+ cells, the lysosomal localization of Mio-FLAG, as indicated by colocalization with the endosomal marker Rab7, was dramatically reduced in *ragA*^11^ mutant clones (**Fig 2A**). Similarly, the colocalization of the GATOR1 component Nprl3 with Rab7 was greatly diminished in *ragC*^13^ germline clones (**Fig 2B**). To determine if GATOR1 or GATOR2 are required for the lysosomal association of the Rag GTPase, we performed the reciprocal experiments. First, we generated homozygous null mutant clones of the GATOR1 components, *iml1*^13^ and *nprl3*^1^ and examined the localization of the Rag GTPase component Rag C-GFP. We found that Rag C-GFP colocalized with Rab7 in both *iml1*^13^ and *nprl3^1^* mutant germline clones (**Fig 2. C,D)**. To determine the role of the GATOR2 complex in the lysosomal localization of the Rag GTPase, we examined the localization of RagC in *wdr24^1^* null mutant ovaries. We found that in *wdr24^1^* mutant egg chambers, RagA-GFP colocalized with the lysosomal marker Lamp1 in both germline and somatic cells **(Fig2. E,F)**. Taken together, these data indicate, that the recruitment of the Rag GTPase to lysosomes does not require GATOR1 or GATOR2.

**Figure 2.**
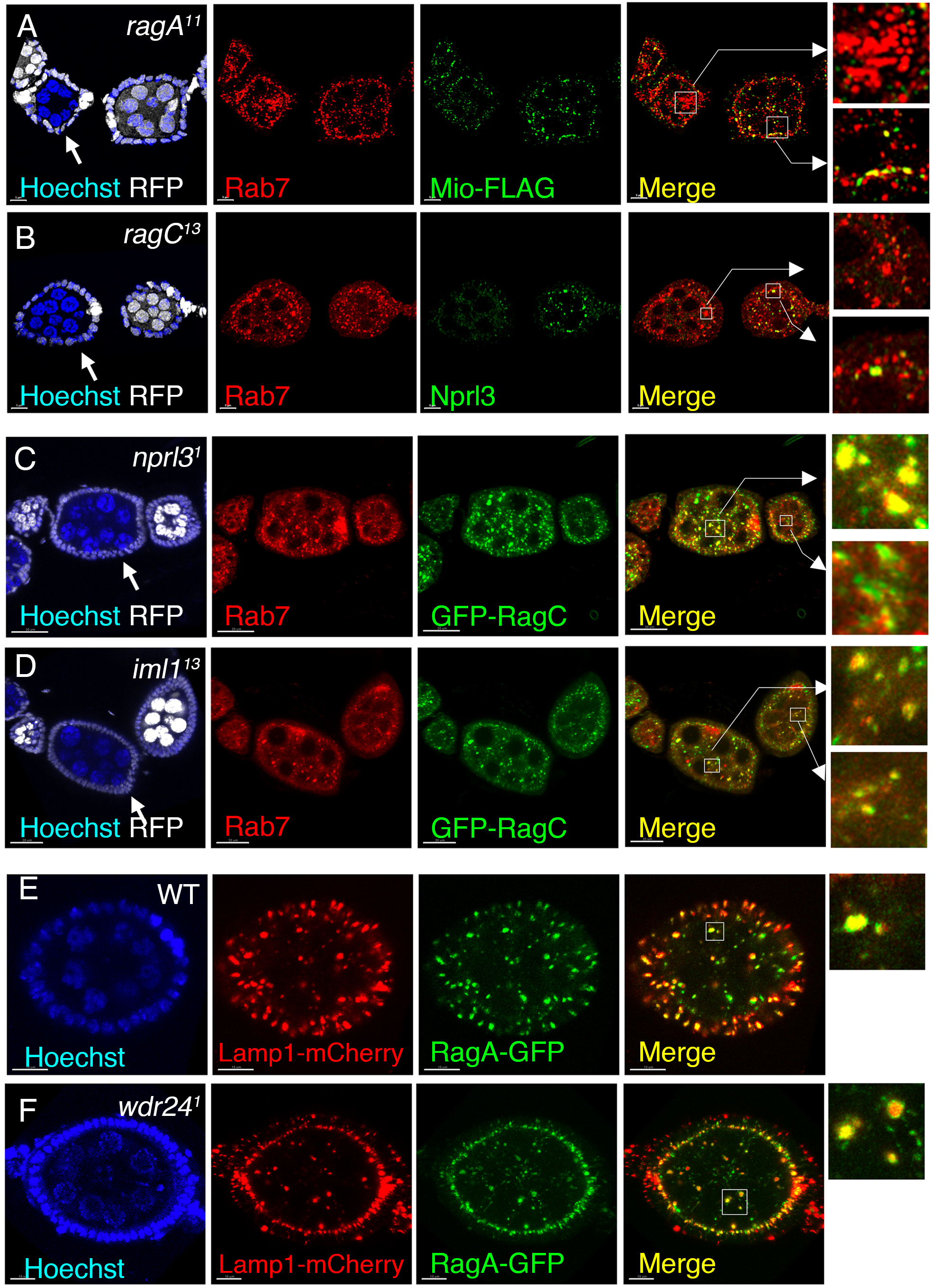
The Rag GTPase localizes GATOR1 and GATOR2 to lysosomes. (A, B) Ovaries containing *ragA*^11^ (A) and *ragC*^13^ (B) homozygous germline clones are marked by the absence of RFP (white, arrow). Mio-FLAG (A, green) and Nprl3 (B, green) were labeled in *ragA*^11^ and *ragC*^13^ mutants, respectively. Lysosomes were marked with anti-Rab7 antibody (Red), and Hoechst 33342 (blue) was used to label DNA. Note that the Mio-FLAG and Nprl3 protein were largely disperse from Rab7 puncta in *ragA*^11^ and *ragC*^13^ mutants respectively (insert A and B). Scale bar: 20 μm. (C, D) *nprl3^1^* (C) and *iml1^13^* (D) homozygous germline clones were marked by the absence of RFP (white, arrow). Rab7 antibody highlights lysosomes while Hoechst 33342 labels DNA. Note that RagC colocalized with Rab7 in *nprl3^1^* and *iml1^13^* mutants. Scale bar:20 um (E) Live cell imaging of *Drosophila* egg chambers showing that RagA-GFP colocalizes with lysosomal marker Lamp1-mCherry in WT (E) *wdr24^1^*mutant ovaries(F). Scale bar:10 um.

Finally, we examined how GATOR1 affects the localization of GATOR2 to lysosomes, and, in turn, how GATOR2 influences the localization of GATOR1. First, we examined the role of GATOR1 in lysosomal localization of the GATOR2 component Mio-FLAG. We determined that in null mutant clones of *nprl3^1^* Mio-FLAG failed to localize to lysosomes (**Fig. 3A**). Similarly, in homozygous germline clones of the null allele *mio^KO2^*, the lysosomal localization of GATOR1 components Iml1 and Nprl2 were dramatically diminished, as determined by their colocalization with Rab7 (**Fig. 3B-C**). Additionally, we found that in *wdr24^1^* mutant egg chambers, the colocalization of GATOR1 components Iml1-V5 and Nprl3, with the lysosomal marker Lamp1-mCherry, were strongly diminished in both germline derived nurse cells and somatically derived follicle cells demonstrating that the requirement for GATOR2 to recruit and/or retain GATOR1 on lysosomes is not confined to the germline (**Fig. 3D-G**). To confirm that in *wdr24^1^* mutant ovaries a smaller fraction of GATOR1 was associated with lysosomes, we performed lysosomal immunoprecipitations (IPs) using Lyso-3XmCherry and determined the ratio of cytoplasmic versus lysosomal Nprl3 in wildtype and *wdr24^1^* mutant ovaries. We calculated that *wdr24^1^* mutants have an approximately 3-fold reduction in the ratio of lysosomal to cytoplasmic Nprl3 (**Fig 3. H,I)**. Thus, similar to what is observed in mammalian cells, GATOR1 and GATOR2 are mutually dependent for their recruitment and/or retention on lysosomes ^29^. However, we note that the low levels of GATOR1 on lysosomes in GATOR2 mutants may be due to the absence of GATOR2 activity, which renders GATOR1 constitutively active. As a result, it is possible the majority of RagA-GTP is converted to RagA-GDP. Because GATOR1 has a high affinity for RagA-GTP, but not for RagA-GDP, its association with lysosomes may be reduced ^39,41,42^.

**Figure 3.**
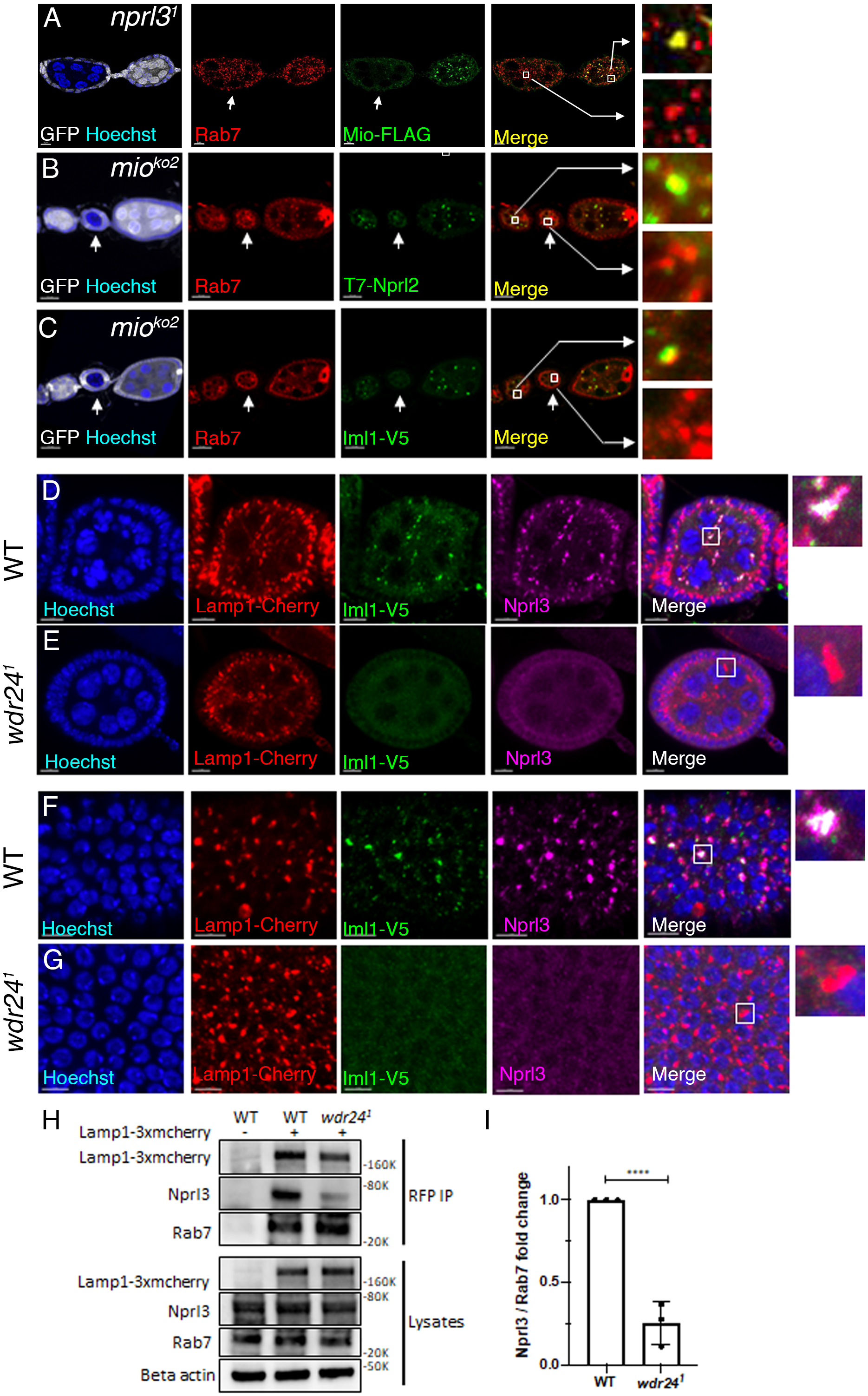
Localization of GATOR1 and GATOR2 to lysosomes is interdependent. (A) *nprl3^1^* homozygous germline clones are marked by the absence of GFP(white, arrow). Lysosomes are labeled with anti-Rab7 antibody (Red) and Hoechst 33342 labels DNA. Note that the Mio-FLAG protein largely disperses from Rab7 puncta in *nprl3^1^* mutants (insert A). (Scale bar: 20 um). (B, C) *mio^ko2^* homozygous germline clones are marked by the absence of GFP (white, arrow). anti-T7 and anti-V5 label Nprl2 and Iml1, respectively (green). Rab7 (red) antibody highlights lysosomes and autolysosomes and Hoechst 33342 labels DNA. In *mio^ko2^* mutant, Nprl2 and Iml1 proteins are largely disperse from Rab7 positive puncta (Scale bar: 20 um). (D-G) Whole egg chambers (D,E) and follicle cells (F,G) from wild-type and *wdr24^1^* females carrying Iml1-v5-GFP^11^ knock-in and Lamp1-3xmcherry (Red) genomic insertions, co-stained with anti-Nprl3 (Magenta) and anti-v5 for Iml1(Green). In *wdr24^1^* mutants, Iml1 and Nprl3 are largely disperse from Lamp1-3xmcherry puncta in both germline and somatic follicle cells (F, G insert).(Scale bar:7 um for D,E. 5um for F,G). (H) Ovaries from wild-type and *wdr24^1^* females carrying Lamp1-3xmcherry genomic insertions. Ovarian lysosomes from each genotype were immunoprecipitated with anti-RFP beads. Lysates (input) and immunoprecipitates (IP) were detected by Western blot using RFP for Lamp1, Nprl3, Rab7 and Beta-actin antibodies. (I) Quantification of three independent experiments from (H). Error bars represent the standard deviation (SD). ****: p<0.0001.

Taken together these data demonstrate that, even though *Drosophila* does not have proteins homologous to those that make up KICSTOR, many of the regulatory hierarchies that control the recruitment of GATOR2, GATOR1, and the Rag GTPase to lysosomes are conserved between mammals and *Drosophila*^38,39^. Finally, we found that unlike other components of GATOR2, Wdr59 was not required for the localization of GATOR1 to lysosomes or autolysosomes **(Supplemental Fig. 2)**. These data are consistent with our previous results that Wdr59 is not required for the lysosomal localization of the other GATOR2 complex components Wdr24 and Mio ^22^. Moreover, they are consistent with Wdr59/SEA3 having a unique function within GATOR2/SEACAT ^22,30–32^.

### GATOR2 promotes the GAP inactive mode of GATOR1

How GATOR2 inhibits the GAP activity of the GATOR1 complex remains poorly understood. Multiple studies indicate that the GATOR2/SEACAT complex does not act as an active site inhibitor of GATOR1 GAP activity but likely functions as an allosteric inhibitor ^39,41,42^. In mammalian cells, structural and biochemical analysis indicate that GATOR1 binds to the Rag GTPase in either an activated (GAP active) or inhibited (GAP inactive) configuration ^42,43^. In the inhibited configuration, the DEPDC5/Iml1 subunit of GATOR1 has a high affinity contact with RagA of the Rag GTPase, which inhibits the GATOR1-mediated hydrolysis of RagA^GTP^. In contrast, in the more transient GATOR1 GAP active form, there is a weak interaction between the Nprl2 and the Rag GTPase through its target RagA ^43^. A similar structure for the GAP-active form of SEACIT, the counterpart to GATOR1, occurs in yeast, but there is no evidence for a GAP-inactive form of SEACIT ^26,39^.

*Drosophila* contains domains homologous to mammalian Iml1/DEPDC5 and RagC domains that are critical for the formation of the GAP-inactive form of GATOR1. Therefore, we examined if the GATOR2 complex impacts the transition of GATOR1 from a GAP inactive to a GAP active form in *Drosophila*. As an entry point for these studies, we used our CRISPR knock-in GATOR1 proteins. First, we performed immunoprecipitations of a tagged Iml1-V5 protein, followed by mass spectrometry, using lysates from both wildtype and *wdr24^1^*mutant ovaries (**Fig 4A, B)**. As expected, in wildtype ovaries, Iml1-V5 co-immunoprecipitated with components of both the GATOR1 (Nprl2/Nprl3) and GATOR2 complexes, although SEC13, a GATOR2 component, was not co-immunoprecipitated (**Fig. 4A**). In contrast, immunoprecipitation of Iml1-V5 from *wdr24^1^* mutant ovaries failed to co-immunoprecipitate, Nprl2 or Nprl3, as well as most members of the GATOR2 complex. To confirm these results, we performed immunoprecipitations of Iml1-V5 followed by western blot analysis for Iml1-V5 and Nprl3 in both wildtype and *wdr24^1^*mutant ovaries. Consistent with the mass spectrometry, the ratio of co-immunoprecipitated Nprl3 to Iml1-V5 was sharply reduced in *wdr24^1^*mutants relative to wildtype, while the overall levels of Nprl3 remained largely unchanged (**Fig 4C, D)**. Therefore, in the absence of GATOR2, where the GATOR1 complex is predicted to be constitutively active, Iml1 shows a reduced affinity for the other two components of the GATOR1 complex, Nprl2, which contains the GAP catalytic site, and Nprl3. Notably, we found that in WDR24-KO HeLa cells, the association of DEPDC5/Iml1 with NPRL2 is diminished suggesting this role of GATOR2 may be conserved in mammals **(Supplemental Fig. 3A, B)**.

**Figure 4.**
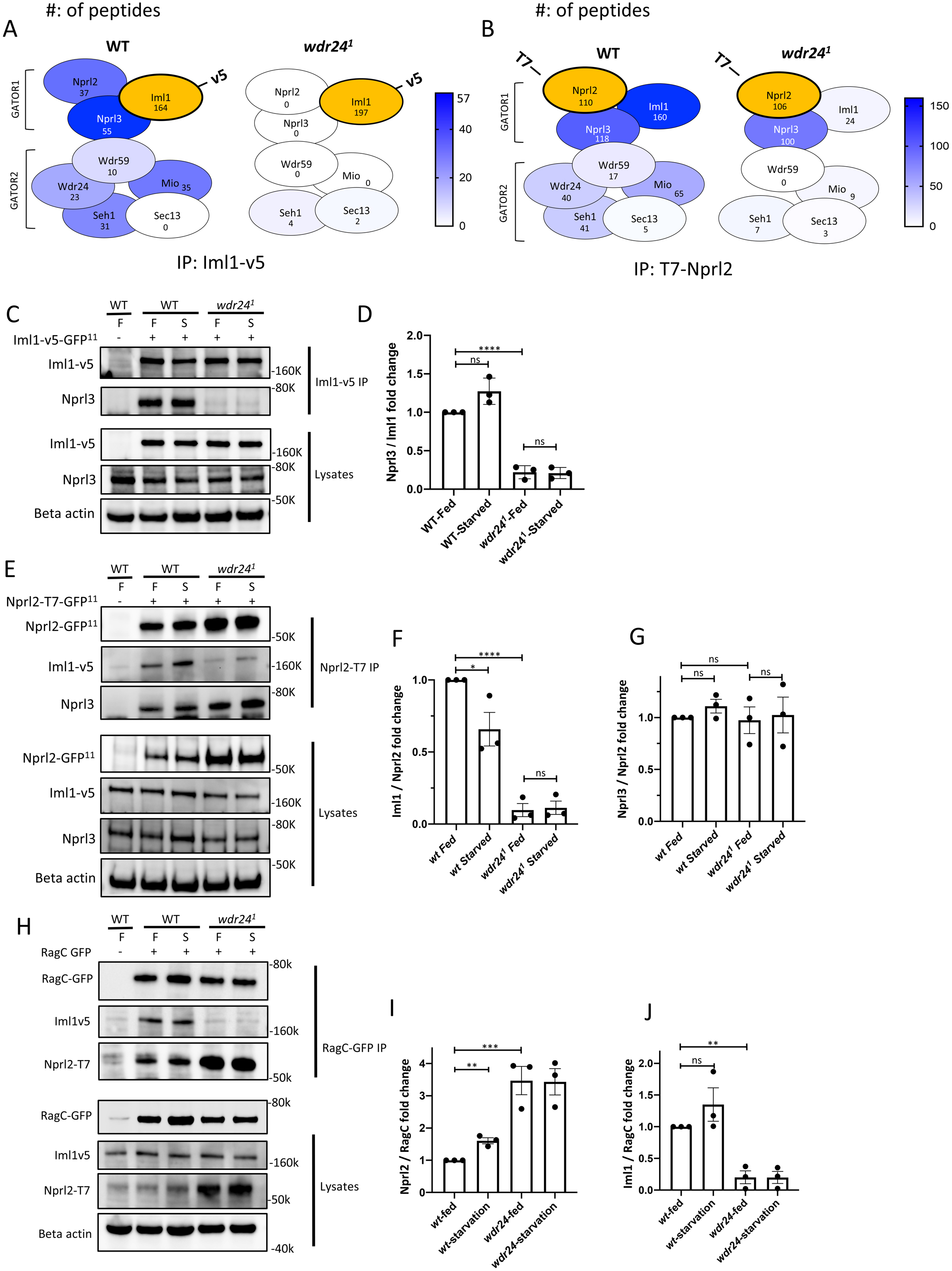
GATOR2 regulates the association of Nprl2/Nprl3 with Iml1 and the Rag GTPase. (A, B) Heat map cartoons summarizing peptide counts form three independent mass spectrometric analyses of anti-v5 (A, Iml1-v5), anti-T7 (B, Nprl2-T7) immunoprecipitates from ovarian lysates of WT and *wdr24^1^* mutant females. GATOR components are color-coded according to the total peptide counts. (C, D) Western blot confirming the association of Iml1 with Nprl3 is decreased in *wdr24^1^* mutant ovaries. Ovaries were dissected from fed and starved, WT, *wdr24^1^* females carrying the Iml1-v5-3xGFP^11^ knockin. Ovarian lysates (input) from each genotype and condition were IPed with anti-v5 antibodies and detected by western blot using antibodies against v5, Nprl3 and Beta-actin. (D) Quantification of Nprl3 levels relative to Iml1 by western blot from (C) (N = 3). (E-G) Immunoblot confirming the association of Nprl2 with Iml1, but not Nprl3, is decreased in *wdr24^1^* mutant ovaries. Ovaries were dissected from fed and starved, WT and *wdr24^1^* females carrying Iml1-v5-3xGFP^11^ and Nprl2-5xT7-3xGFP^11^ knockins. Ovarian lysates (input) from each genotype and condition were IPed with anti-T7 antibodies. Ovarian lysates and IPs were detected by western blot using antibodies against T7 for Nprl2, v5 for Iml1, Nprl3 and Beta-actin. (F) Quantification of Iml1 levels relative to Nprl2 by western blot from (E) (N = 3). (G) Quantification of Nprl3 levels relative to Nprl2 by Western blot from (E) (N = 3). (H-J) GATOR2 inhibits the association of Nprl2 with the Rag GTPase. (H) Ovaries were dissected from fed and starved, WT and *wdr24^1^* females carrying Iml1-v5-3xGFP^11^, Nprl2-5xT7-3xGFP^11^ knockins and a RagC-GFP genomic insertion. Ovarian lysates were IPed with anti-GFP antibodies. Ovarian lysates and IPs were detected by Western blot using antibodies against T7 for Nprl2, v5 for Iml1, GFP for RagC and Beta-actin. (I) Quantification of Nprl2 levels relative to RagC by Western blot from (H) (N = 3). (J) Quantification of Iml1 levels relative to RagC by Western blot from (H) (N = 3).

To confirm our results, we immunoprecipitated a second endogenously tagged component of GATOR1, Nprl2-T7, and analyzed the precipitate by mass spectrometry. As observed with Iml1-V5, in wildtype ovaries, Nprl2-T7 co-immunoprecipitated the other two components of the GATOR1 complex, Nprl3 and Iml1, as well as subunits of the GATOR2 complex (**Fig. 4B**). However, in *wdr24^1^* mutant ovaries, while Nprl2-T7 robustly co-immunoprecipitated Nprl3, its association with Iml1 and GATOR2 complex proteins was sharply decreased. To verify these findings, we performed immunoprecipitations of Nprl2-GFP^11^ followed by western blot analysis for Iml1-V5 and Nprl3 in lysates from wildtype and *wdr24^1^* mutant ovaries. These data confirmed that the association between Nprl2 and Nprl3 was not notably altered in *wdr24^1^* mutants compared to wildtype, as determined by the ratio of Nprl2/Nprl3, however, the ratio of Nprl2/Iml1 was substantially decreased (**Fig. 4E-G**). These results suggest that the GATOR2 complex, which opposes GATOR1 activity, promotes a high affinity association of Iml1 with the Nprl2/Nprl3 heterodimer. Notably, we also observed that the levels of Nprl2 protein were increased in the *wdr24^1^* mutant background **(Supplemental Fig. 4)** ^44^.

To investigate how GATOR2 impacts the association of GATOR1 components with the Rag GTPase, we performed immunoprecipitations using a genomically tagged RagC-GFP protein, followed by western blotting for the GATOR1 components Iml1-V5 and Nprl2-T7. As expected, in wild-type ovaries, Rag-C-GFP co-immunoprecipitated with both Iml1-V5 and Nprl2-T7, confirming that the Rag GTPase associates with the GATOR1 complex under both fed and starved conditions **(Fig4. H-J)**. Additionally, as we previously reported, the association of the Rag GTPase with Nprl2 increased under starvation conditions in *wdr24^1^* mutant ovaries, where GATOR1 is constitutively active ^22^. However, consistent with the differential behavior of the Iml1 component of GATOR1 in GATOR2 mutants, the ability of Rag-C-GFP to co-immunoprecipitate Iml1-V5 protein was significantly reduced in *wdr24^1^* mutant ovaries **(Fig4. H-J)**. Thus, in the absence of its inhibitor GATOR2, the catalytic component of GATOR1, Nprl2, shows an increased affinity for its target, the Rag GTPase, while Iml1 exhibits a decreased affinity. Taken together, our data support the hypothesis that the GATOR2 complex promotes the association of the Nprl2/Nprl3 heterodimer with Iml1/DEPDC5 and is critical for maintaining the GATOR1 complex in an inhibited (GAP-inactive) state.

### Starvation decouples the lysosomal recruitment of GATOR1 and GATOR2

To further explore how GATOR2 regulates GATOR1, we used fluorescence recovery after photobleaching (FRAP) to analyze the dynamic behavior of components of the Rag GTPase, GATOR1, and GATOR2 under fed and starved conditions. FRAP measures the rate of exchange between protein populations and allows for the determination of the mobile fraction of a protein that is free to exchange between pools ^45,46^. Previously, we developed a method to perform lysosomal FRAP on whole cultured *Drosophila* ovaries to examine the exchange of proteins between the lysosome and the cytoplasm ^47^. Here we used GFP or a split GFP system, in which rGFP is reconstituted from genomically tagged proteins, with GFP^1–10^ specifically expressed in the female germline (Fig. 1) ^22^. As proof of principle, we first performed lysosomal FRAP to define the lysosomal exchange of an endogenously tagged RagA-GFP protein. Previous work in mammalian cells, reported that nutrients control the association of the Rag GTPase with the lysosome, with the nutrient dependent GTP-loading of RagA increasing the lysosomal-cytoplasmic exchange of the complex^48^.

To examine the behavior of the Rag GTPase in *Drosophila*, we photobleached individual RagA-GFP–labeled lysosomes in stage 6 egg chambers and measured fluorescence recovery under fed and starved conditions (**Fig. 5A,B**). We found that, as is observed in mammalian cells, the Rag GTPase traffics between the cytoplasm and the lysosomal surface in response to nutrient conditions. Under nutrient-replete (fed) conditions, RagA-GFP recovered approximately 30% of its initial fluorescence intensity, corresponding to a mobile fraction of ∼20% (**Fig 5A-C**). In contrast, in ovaries from starved females, the recovery of RagA-GFP fluorescence intensity was reduced twofold. These data indicate reduced recruitment and/or exchange of RagA-GFP to lysosomes under nutrient limitation, consistent with findings in mammalian tissue culture cells^48^. In *wdr24^1^* mutant ovaries, the GATOR1 complex is constitutively active, favoring the Rag GTPase being in the inactive RagA-GDP-bound state ^17,19,23^. Thus, as expected, lysosomal FRAP performed on ovaries from fed females from the GATOR2 mutant *wdr24^1^* mimicked starvation conditions, showing a substantial reduction in fluorescence recovery and mobile fraction of RagA-GFP compared to wild-type fed ovaries (**Fig. 5A-C**).

**Fig 5.**
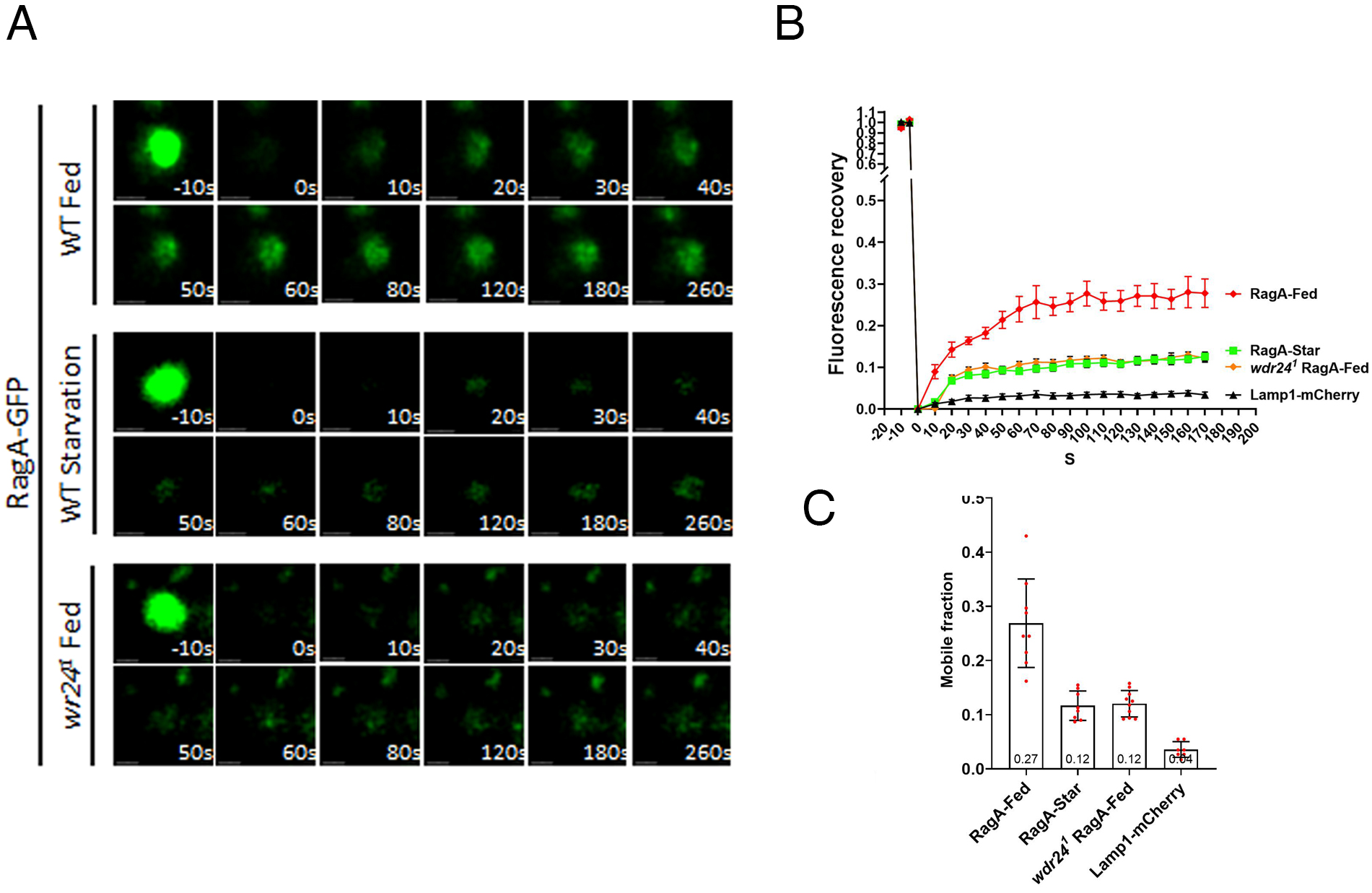
Lysosomal recruitment of RagA to lysosomes is nutrient dependent. (A) Time point images of FRAP experiments in Drosophila ovarian tissue of RagA-GFP under fed, starvation and fed condition in *wdr24^1^*mutant background. (B) Fluorescence recovery versus time curves calculated from (A). (C) Mobile fraction for Rag-GFP proteins.

Next we defined the lysosomal cytoplasmic dynamics of components of both GATOR1 and GATOR2. First, we performed lysosomal FRAP on two GATOR1 components, Nprl2 and Iml1. Under nutrient-replete conditions, the recovery kinetics of Nprl2-rGFP closely mirrored those of RagA-GFP, with approximately 30% recovery of its fluorescent intensity (**Fig 6A, C).** Similarly, Iml1-rGFP exhibited comparable mobility, although with a slightly higher mobile fraction, recovering approximately 40% of its initial fluorescence (**Fig. 6B, C**). As observed with RagA-GFP, both Nprl2-rGFP and Iml1-rGFP, show a marked decrease in lysosomal recruitment upon starvation, with their mobile fractions reduced by more than two-fold (**Fig. 6A-D**). We note that the slight increase in the mobile fraction of Iml1-rGFP likely indicates that Iml1 is somewhat more dynamic within the GATOR1 complex under fed conditions than Nprl2-rGFP. Thus, as is observed with the Rag GTPase, in nutrient replete conditions a fraction of GATOR1 freely exchanges between the cytoplasm and the lysosomal surface and this exchange is restricted upon nutrient limitation.

**Figure 6.**
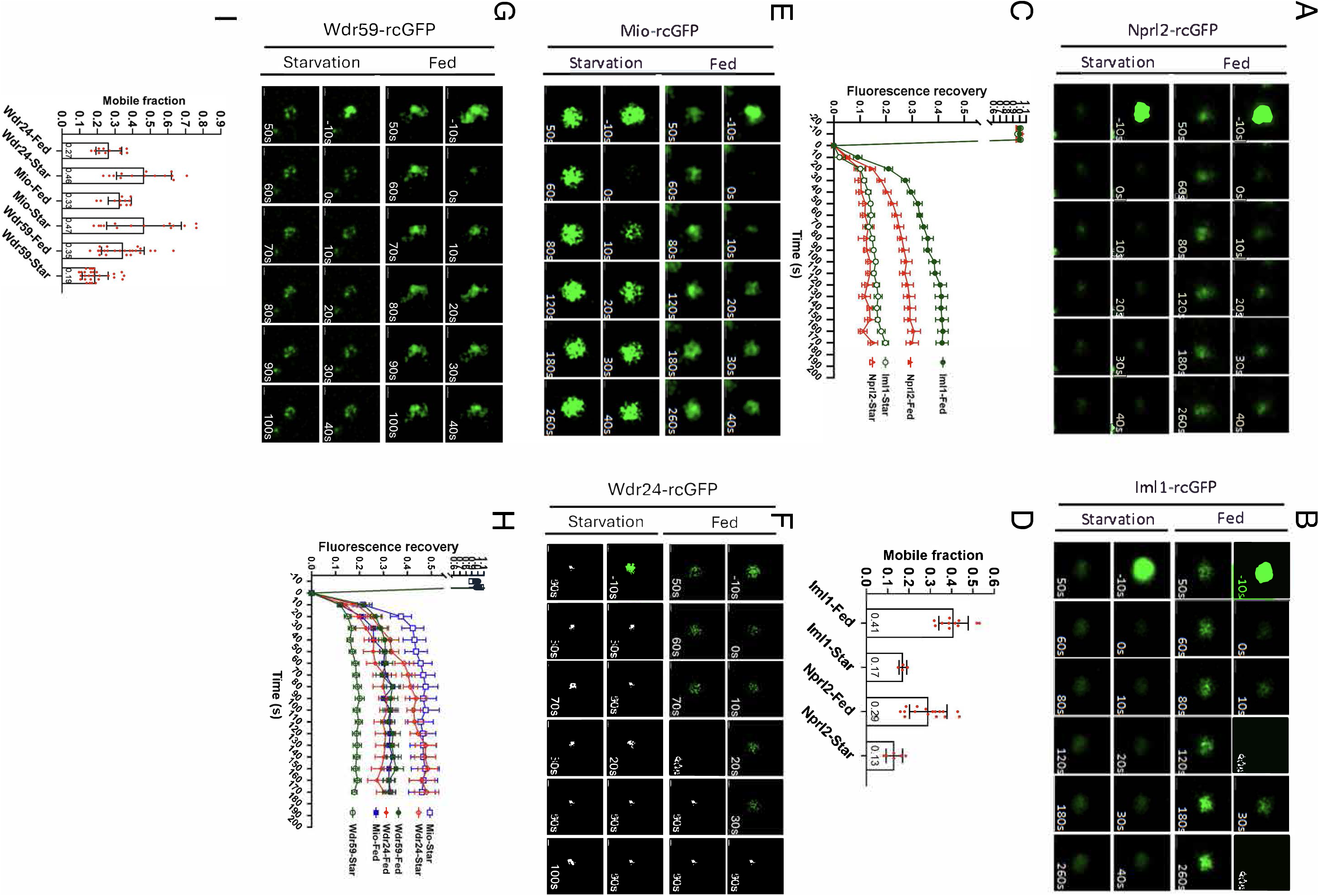
Lysosomal recruitment of GATOR2 mirrors GATOR1 and the Rag GTPase in fed but not starved conditions. (A,B,E,F) Time point images of FRAP experiments in Drosophila ovarian tissue of Nprl2-rGFP (A), Iml1-rGFP (B), Mio-rGFP ( D) and Wdr59-rGFP (F). In fed and starved conditions. The 0s image represents photobleaching. Scale bar: 1 µm. (C,G) Fluorescence recovery versus time curves calculated from (A, B, E, F). Curves were calculated from at least 10 lysosomes from 10 different ovaries for each treatment and genotype. Error bars represent standard error. (D,H) Mobile fraction for tagged proteins under nutrient-replete and nutrient-deplete conditions.

Finally, we examined the dynamic behavior of the core GATOR2 components Mio and Wdr24. Lysosomal FRAP of Mio-rGFP and Wdr24-rGFP revealed that in nutrient-replete conditions, the recovery kinetics of these GATOR2 components resembled that of Iml1-rGFP, Nprl2-rGFP, and RagA-rGFP, with a mobile fraction of approximately 30% (**Fig. 6E,F,H,I**). In contrast to GATOR1 and RagA, however, starvation increased the mobile fraction of Mio-rGFP and Wdr24-rGFP rather than decreases the mobile fraction. Thus, in starvation conditions, where the GATOR2 complex no longer opposes the GATOR1 GAP activity toward RagA, the Wdr24 and Mio components of GATOR2 more rapidly exchange with the lysosome. Recent studies report that GATOR1 and GATOR2, form a supercomplex that traffics between lysosomes and the cytoplasm ^26,38,39^. Taken together our data are consistent with GATOR2 being present in a supercomplex with GATOR1 in fed conditions but having a more transient association in starvation conditions, at a time when GATOR2 is not actively inhibiting the GAP activity of GATOR1.

### The lysosomal-cytoplasmic exchange of Wdr59 mirrors GATOR1 not GATOR2

Wdr59 was originally classified as a GATOR2 subunit based on physical association and functional studies in mammalian and *Drosophila* tissue-culture systems ^17,47,49^. However, recent in vivo studies determined that in many *Drosophila* tissues, including the ovary, Wdr59 functions to oppose GATOR2 activity, with null mutations in *wdr59* resulting in increased TORC1 activity due to the constitutive inhibition of GATOR1 activity by GATOR2^22^. Notably, structural analyses have shown that WDR59/SEA3 is a component of both the GATOR1/SEACIT and GATOR2/SEACAT complexes ^28^. Specifically, WDR59/SEA3’s N-terminal region associates with GATOR1, whereas its C-terminal region contributes to the architecture of GATOR2/SEACIT ^26,39^. Yet, the function of Wdr59 remains mysterious.

Using lysosomal FRAP we determined that in fed conditions, Wdr59-GFP has lysosomal recruitment dynamics similar to other components of GATOR2 and GATOR1 (**Figure 6G-I**). Again, this is consistent with Wdr59 being present in a supercomplex with GATOR1 and GATOR2 in nutrient-replete conditions as suggested by work in yeast and mammalian cells ^26,38,39^. However, upon exposure to nutrient limitation, Wdr59-GFP dynamics closely mirror those of GATOR1, with the Wdr59-GFP fluorescence recovery and mobile fraction being reduced in response to starvation rather than increased as we observe with the GATOR2 components Mio and Wdr24. These data indicate that Wdr59 becomes at least partially disengaged from the GATOR2 complex in response to starvation.

We were interested in determining how mutations in *wdr59*, which result in the constitutive activation of the GATOR2 complex and thus the constitutive inhibition of GATOR1, impact the lysosomal cytoplasmic dynamics of both GATOR1 and GATOR2 ^22^. We predicted that if lysosomal recruitment reflects GATOR1 activity thanGATOR1 would have similar dynamics in fed and starved conditions in in *wdr59* mutants. To test this hypothesis, we performed lysosomal FRAP of Iml1-rGFP (GATOR1) and Mio-rGFP (GATOR2) in *wdr59^1^* null mutant ovaries. First we performed FRAP on the GATOR1 component Iml1-rGFP in fed and starved conditions in *wdr59^1^* mutant ovaries. We found that, as predicted, the fluorescent recover rate and mobile fractions were nearly identical in both conditions with no diminished recovery in response to starvation as is observed for Iml1-rGFP in a wildtype background (**Fig. 7A-D**). These data support our previous finding that in *Drosophila*, *wdr59^1^*mutants have a minimal response to nutrient stress with *wdr59* mutants only slightly downregulating TORC1 activity in response to starvation^22^. Finally, we performed FRAP analysis on Mio-rGFP from *wdr59^1^* mutant ovaries cultured in fed and starved conditions. In fed conditions the lysosomal recruitment of Mio-rGFP mirrored that of Iml1-rGFP in the *wdr59^1^* background. These data are consistent with the GATOR1 and GATOR2 complex forming a supercomplex or alternatively, their recruitment to lysosomes being regulated by the same rate limiting step, even in the absence of the Wdr59 component of GATOR2. However, to our surprise, in starvation conditions, the fluorescent recovery of Mio decreased in *wdr59^1^* mutants. Why the recruitment of Mio to lysosomes decreased in the absence of Wdr59 is currently unclear.

**Figure 7.**
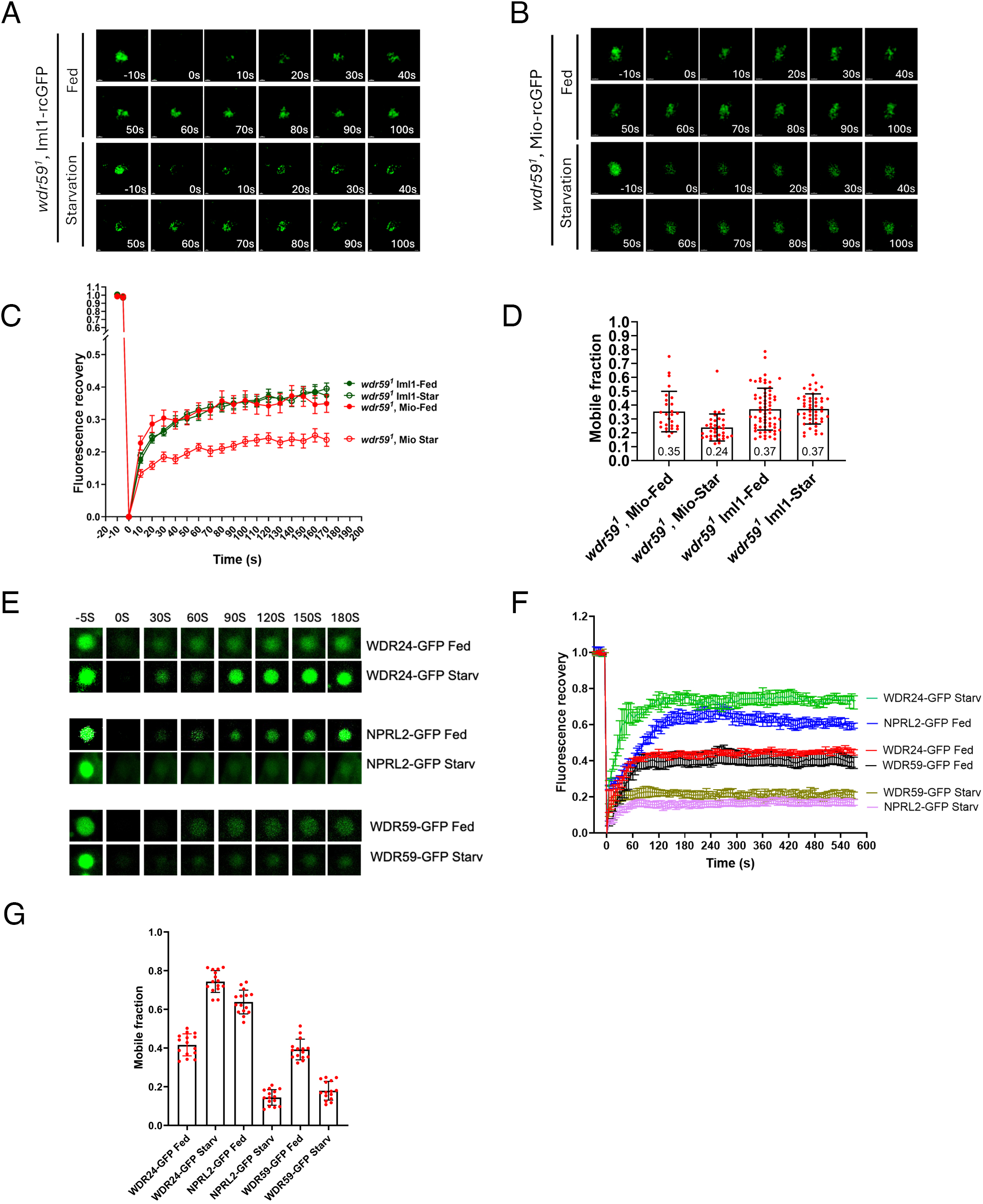
Wdr59 regulates the recruitment of GATOR1 and GATOR2 to lysosomes. (A,B) Time point images of FRAP experiments in *wdr59^1^* mutant ovaries of Iml1-rGFP (A) and Mio-rGFP (B). (C) Fluorescence recovery versus time curves for Iml1-rGFP and Mio-rGFP in *wdr59^1^* mutant ovaries in fed and starved conditions. Curves were calculated from at least 10 lysosomes from 10 different ovaries for each treatment and genotype. Error bars represent standard error. (D) Mobile fraction for tagged proteins shown in (C) under fed and starved conditions. (E) Time point images of FRAP experiments WDR24-GFP, NPRL2-GFP and WDR59-GFP in fed and starved conditions in mammalian HeLa cells. (F) Fluorescence recovery versus time curves for WDR24-GFP, NPRL2-GFP and WDR59-GFP in fed and starved conditions in mammalian HeLa cells. (G) Mobile fraction for tagged proteins shown in (E) under nutrient-replete and nutrient-depleted conditions.

**Figure 8.**
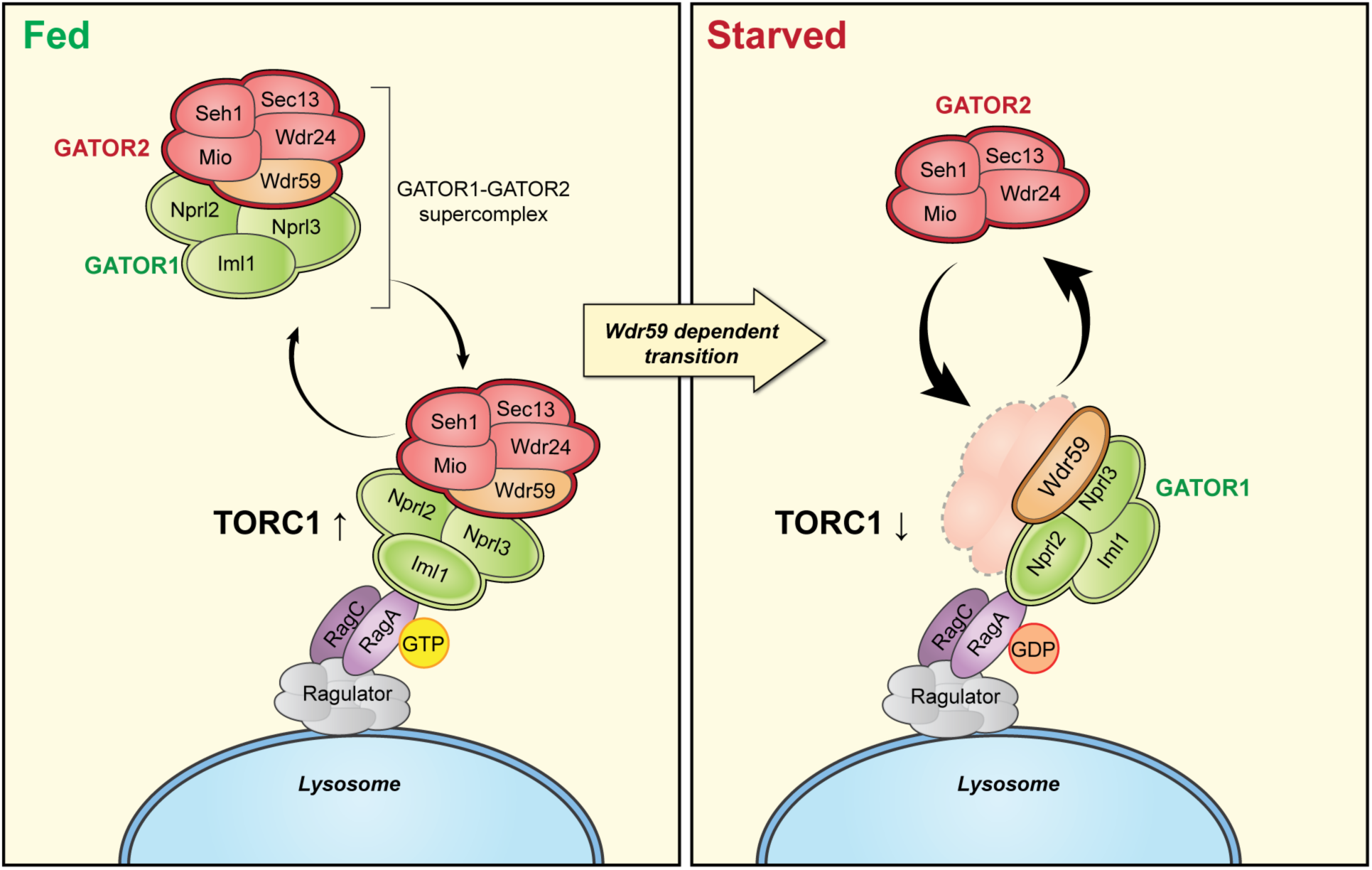
GATOR2 regulates the recruitment of GATOR1. Under nutrient-rich conditions, GAP-inactive GATOR1 forms a supercomplex with GATOR2 that transiently associates with the Rag GTPase, allowing the supercomplex to cycle on and off lysosomes. In response to starvation, the GATOR1-GATOR2 supercomplex dissociates, enabling GATOR1 to adopt a GAP-active state in which Nprl2, the catalytic subunit of GATOR1, acts on the Rag GTPase RagA to inhibit its activity. Wdr59 facilitates the transition of GATOR1 from the GAP-inactive to the GAP-active state. Notably, the lysosomal dynamics of Wdr59 resemble those of GATOR1 rather than GATOR2.

A final question was whether our observations in *Drosophila* on the role of the GATOR2 complex in regulating the lysosomal recruitment and activity of GATOR1 is conserved in mammals. To address this, we examined lysosomal recruitment of the GATOR2 components WDR24-GFP and WDR59-GFP, and the GATOR1 component NPRL2-GFP, using lysosomal FRAP analysis in transiently transfected HeLa cells ^47^. Overall, the trends we observed closely mirrored those in *Drosophila*. Starvation decreased lysosomal recruitment of the GATOR1 component NPRL2-GFP, while increasing the lysosomal recruitment of the GATOR2 component WDR24-GFP (**Fig. 7G-I**). Moreover, WDR59 behaved more like the GATOR1 complex, showing a marked reduction in lysosomal recruitment under starvation conditions. These data support the model that as is observed in *Drosophila*, starvation promotes the differential recruitment of GATOR1 and GATOR2 to lysosomes.

## DISCUSSION

Here we present a molecular genetic resource that provides a platform to dissect TORC1 regulation and its upstream control by the GATOR–Rag GTPase network in Drosophila. By genomically tagging pathway components at their endogenous loci, we have created reagents that allow for the biochemical characterization as well as visualization of these pathway components under native expression levels. These reagents now make it possible to follow the activity, physical association and localization of GATOR2, GATOR1, and Rag GTPase components with high spatial and temporal resolution across diverse cell types of *Drosophila*. This resource provides a versatile toolkit that will facilitate mechanistic and physiological studies of TORC1 regulation in vivo. As proof of principle, we have used these reagents to demonstrate that many of the regulatory principles that govern the GATOR-Rag GTPase signaling pathways in mammals are conserved in *Drosophila*. Moreover, our studies reveal a role for the GATOR2 complex in ensuring that the GATOR1 complex is retained in an inhibited configuration in nutrient replete conditions. Finally, our data support the model that nutrient limitation decouples the lysosomal recruitment of GATOR1 and GATOR2 allowing for active form of GATOR1 to engage the Rag GTPase on lysosomes.

### A toolbox for the study of TORC1 regulation

While components of the GATOR signaling pathway are broadly conserved between mammals and *Drosophila*, there are several species-specific differences in regulatory inputs and complex composition that reflect evolutionary adaptations in nutrient sensing ^9,22,30^. In both organisms, GATOR1 inhibits TORC1 activity by acting as a GTPase-activating protein (GAP) for the RagA component of the Rag GTPase, while GATOR2 relieves this inhibition by suppressing GATOR1 function via an unknown mechanism ^11–13,19^. However, several upstream regulators present in mammals, including the multiprotein KICSTOR complex, which tethers GATOR1 to lysosomes, and CASTOR1, which interacts with GATOR2 to relay information about intracellular arginine, are not conserved in *Drosophila* ^31,40^. The apparent absence of orthologs or functional equivalents for these complexes raised the possibility that amino acid sensing through GATOR2 may utilize an alternative mechanism in *Drosophila*.

To address this possibility, we used our genomically tagged proteins, to examine whether the mechanistic principles governing GATOR–Rag GTPase regulation in mammals are conserved in *Drosophila*. Using immunolocalization of tagged proteins in combination with loss of function genetics we determined that, as is observed in mammalian cells, in *Drosophila*, the Rag GTPase recruits GATOR1 and GATOR2 to the lysosomal membrane, while GATOR1 and GATOR2 are interdependent for their lysosomal localization ^26,38,48^. However, we note that the limited lysosomal localization of GATOR1 in GATOR2 mutants may reflect the nucleotide status of the RagA component of the Rag GTPase in these genetic backgrounds. ^18^ ^26,39,41,42^. GATOR1 has increased affinity to RagA-GTP relative to RagA-GDP^17,29,39^. In GATOR2 mutants, we predict that the preponderance of lysosomal RagA is GDP bound resulting in the release of GATOR1 from the lysosome. Taken together, our findings indicate that the core requirements for recruitment of GATOR1 and GATOR2 in *Drosophila* resembles those found in mammals with the exception of the GATOR2 component Wdr59 which is not required for the lysosomal association of GATOR1 or GATOR2 in the *Drosophila* ovary ^22^.

### GATOR2 promotes the GAP inactive form of GATOR1

Genetic analysis in multiple organisms has found that all three components of GATOR1(Nprl2 Nprl3, and Iml1/DEPDC5) are required for full GAP activity ^16,17,19,21,50,51^. Structural and biochemical studies indicate that in mammals, GATOR1 exists in two distinct conformational states^42^. When GATOR1 is active, the catalytically active subunit Nprl2 associates with its target RagA, thereby enabling GAP activity^42^. When GATOR1 is in the GAP-inactive form, there is a high affinity association between RagA and catalytically inactive Iml1/DEPDC5 via a domain in Iml1/DEPDC5 called the critical strip, which is conserved in *Drosophila* but not in yeast^42^. This interaction is proposed to stabilize the GATOR1 GAP inactive configuration, thus preventing GATOR1-mediated GTP hydrolysis of RagA-GTP ^41–43^. Our data support a role for GATOR2 in stabilizing the GATOR1complex in the GAP inactive conformation. In mutants lacking the GATOR2 component Wdr24, Iml1/DEPDC5 shows reduced association with the Rag GTPase, while the catalytically active Nprl2 exhibits increased association. Moreover, we find that GATOR2 promotes the association of Iml1/DEPDC5 with the Nprl2/Nprl3 heterodimer: this interaction is markedly reduced in *wdr24* mutants, whereas the Nprl2-Nprl3 association remains intact. Based on these observations we propose a model in which GATOR2 inhibits GATOR1 by reinforcing the GATOR1 inactive state in which Iml1/DEPDC5 has a high affinity association with the Rag GTPase and Iml1/DEPDC5 stably associates with the Nprl2/Nprl3 heterodimer.

### Starvation decouples the lysosomal-cytoplasmic exchange of GATOR1 and GATOR2

Early work on the SEA complex reported that components of GATOR1/SEACIT (DEPDC5/Iml1) and GATOR2/SEACAT (Mio/Sea4) dynamically associate with the vacuole ^14^. Using live cell imagining of genomically tagged proteins, we examined the nutrient dependent recruitment of GATOR1 and GATOR2 to lysosomes in the *Drosophila* ovary. We found that under nutrient-replete conditions, both GATOR1 and GATOR2 are recruited to lysosomes with similar dynamics which is consistent with biochemical data from yeast and mammals suggesting that GATOR1 and GATOR2 traffic from the cytoplasm to the lysosome as part of single large supercomplex^26,38,39^. In nutrient replete conditions, the recruitment of the GATOR1-GATOR2 supercomplex resembles that of the Rag GTPase ^38^. Previous studies have found that GTP loading of RagA/B increases the lysosomal off rate of Rag GTPases and controls mTORC1 residence time ^48^. The GATOR1 complex prefers to bind the RagA-GTP form of the Rag GTPase ^17,29,39^. These data are consistent with structural analysis from yeast suggesting the inactive (GDP bound) form of GTR1 (RagA/B) is incapable of binding the SEACIT (GATOR1). Thus, it is likely that the nucleotide state of the Rag GTPase is also determining the residence time and cytoplasmic-lysosomal dynamics of the GATOR1-GATOR2 supercomplex, driving the relatively rapid/transient association of the supercomplex in nutrient replete conditions when RagA is more likely to be GTP bound.

However, upon starvation we observed that the recruitment of GATOR1 to lysosomes decreases sharply⁴⁶. This observation is consistent with GATOR1 converting most of the RagA-GTP on lysosomes, to RagA-GDP, which has a reduced affinity for GATOR1 ^48^. In contrast, we find that lysosomal recruitment of the core GATOR2 components Mio and Wdr24 increase under starvation conditions. At least two models could explain this starvation dependent decoupling of GATOR1 and GATOR2 lysosomal recruitment. First, starvation may promote the recruitment or retention of a pool of GATOR2 that exists independently of the GATOR1–GATOR2 supercomplex. However, this would necessitate that starvation increases GATOR2 on lysosomes which is the opposite of several recent reports ^52,53^. Therefore, we favor a second model in which starvation triggers disassembly and/or reorganization of the supercomplex, relieving GATOR2-mediated inhibition of GATOR1 and enabling GATOR1 to adopt its GAP-active conformation. Based on both the present data and our previous work, we further hypothesize that the transition from the GAP-inhibited to GAP-active state of GATOR1 is at least partially facilitated by Wdr59 ^22^. Consistent with this idea, *wdr59* mutants fail to show starvation-dependent changes in GATOR1 lysosomal recruitment. While these results are suggestive, a complete understanding of what regulates the nutrient dependent recruitment of GATOR1 and GATOR2 to lysosomes awaits further analysis ^22^.

In summary we have generated tools to explore the regulation of the GATOR-Rag GTPase-TORC1 signaling pathway at the cellular and subcellular level in multiple tissues and genetic backgrounds. We demonstrate that GATOR2 maintains GATOR1 in a GAP-inactive state. Moreover, we find that the GATOR1-GATOR2 supercomplex is a nutrient-responsive system whose subcomplexes exhibit distinct cytoplasmic-lysosomal dynamics. In the future it will be important to determine the precise mechanism by which GATOR2 impacts the lysosomal recruitment and activity of GATOR1 as well as the precise function of Wdr59 in this process. fraction on the lysosomes. Defining the importance and regulation of this immobile fraction should be an important goal for future studies.

## Acknowledgments

Multiple stocks used in this study were obtained from the Bloomington *Drosophila* Stock Center supported by NIH grant P40OD018537. Information from Flybase was used in this work which is supported by NIH grant P41 HG00739. We thank Alexander Sodt for his insightful comments on the FRAP analysis.

## Author contributions

Conceptualization: Y. Z., C. Y.T. and M.A.L. Methodology: Y.Z., C.-Y.T., and M.A.L. Investigation: S.Y., R. G., S. Y. and C.-Y.T. Funding acquisition: M.A.L and S. Y. Project administration: M.A.L. and C.-Y.T. Supervision: M.A.L. Writing – original draft: Y. Z. and M.A.L. Writing – review & editing: Y. Z., C.-Y.T., R. G. S. Y. and M.A.L.

## MATERIALS AND METHODS

### *Drosophila* Strains and Genetics

The information details of the flies used in this study were listed in the Key resources table. All *Drosophila* stocks were maintained at 25 ° C on JAZZ-mix *Drosophila* food (Fisher Scientific). Experimental animals were collected at 3-5 days and were well-fed with wet yeast. 1xPBS was used for starvation treatments for 12 hours.

### CRISPR design

The CRISPR target sites were designed via flyCRISPR Target Finder Tool on http://flycrispr.molbio.wisc.edu/tools

### gRNA construct

CRISPR gRNA was made by self-annealing of the two oligos and sub-cloned into BbsI site of pU6-BbsI-chiRNA (Addgene 45946) or pCFD4-3xP3DsRed (Addgene 86864).

### Generation of raga, ragC and iml1 knockout alleles

#### ragA^11^

gRNA (TTGACTAGCGTAGCCTCCGAGGG) was cloned into the pCFD4-3xP3DsRed plasmid (addgene 86864). The recombinant DNA was injected into nos-Cas9 bearing flies to make knockout alleles.

#### ragC^13^

Two gRNA were cloned into pCFD4-3xP3DsRed plasmid. The recombinant DNA was injected into nos-Cas9 bearing flies to make knockout alleles.

gRNA1: TTCCAGATCTGGGACTTCCCCGG

gRNA2: GATCGAGGCCAGCCTCGTTAAGG

#### iml1^13^

Two gRNA were cloned into pCFD4-3xP3DsRed plasmid. The recombinant DNA was injected into nos-Cas9 bearing flies to make knockout alleles.

gRNA1: GTGGTCGCAAGGCGAGCGCGTGG

gRNA2: AAGCTCAAGTTTATCCAGTGGGG

### RagA genomic constructs

RagA-3xHA-1xMyc and RagA-1xGFP-3xHA-1xMyc:

The 4.4 kb of RagA genomic sequence, which included a 1.5kb promoter-enhancer region and a 1.3kb 3’ region (including the intron), was amplified by PCR. The DNA encode 3xHA-1myc or GFP-3xHA-1myc were subsequently assembled into RagA genomic DNA by In-Fusion cloning to make c terminal tagged RagA genomic constructs.

### GFP-RagC genomic construct

The 5.3 kb of RagC genomic fragment, which included a 1.8 kb promoter-enhancer region and a 1 kb 3’ region (including intron), was amplified by PCR.The DNA encode GFP fragment was subsequently assembled into RagC genomic DNA by In-Fusion cloning to make N terminal tagged RagC genomic constructs.

### Genomic knock-in

Genomic knock-in were performed by CRISPR knock-in as described previously ^54^. The detailed design was provided upon request.

### Fly injection

gRNA (100 ng/mL) and targeting (250 ng/mL) constructs were co-inject into y[1], sc[1], v[1];; nos-cas9, v, attp2 or y[1], sc[1], v[1]; nos-cas9, v, attp40. Injection services were provided by Rainbow Transgenic Flies or BestGene Inc.

### Western Blot Analysis

*Drosophila* ovaries were homogenized in RIPA buffer containing Complete Protease Inhibitors and Phosphatase Inhibitors (Roche). Antibodies were used at the following concentrations: rabbit anti-P-S6K T398 (1:1000) (Cell Signaling), guinea pig anti-S6K (1:10,000) (PMID: 20444422), guinea pig anti-Nprl3 (1:1000) (^22^), mouse anti-actin (1:10000)(Abcam), rabbit anti-GFP^11^ (1:1000) (Bonopus), rat anti-OLLAS (1:1000) (Thermo Fisher), mouse anti-T7 (1:1000) (Millipore sigma), rabbit anti-GFP (1:1000) (Cell signaling). The band intensity was quantified using Image J analysis tool (NIH).

### Immunofluorescence and Live Imaging

Immunofluorescence was performed as previously^22,47^, using the following antibodies: rabbit anti-GFP^11^ (1:1000) (Bonopus), guinea pig anti-Nprl3 (1:1000), mouse anti-Rab7 (1:100) (Developmental Studies Hybridoma Bank), rabbit anti-GFP (1:1000) (Cell Signaling Technologies), mouse anti-T7 tag (1:1000) (Millipore sigma), rabbit anti-v5 tag (1:1000) (Thermo Fisher Scientific). Anti-rabbit, anti-mouse and anti-guinea pig Alexa Fluor secondary antibodies (Invitrogen) were used at 1:1000. Nuclei were visualized by staining the DNA with Hoechst 33342 (Thermo Fisher, #H3570) or DAPI.

### Clonal Analysis

NosFlp; *mio^ko^,* FRT40 / FRT40, Ubi-GFP females, NosFlp;; *ragA^ko^,* FRT82b / FRT82b, Ubi-RFP females, NosFlp; *ragc^ko^,* FRT42D / FRT42D, Ubi-RFP females, NosFlp;; *nprl3^ko^,* FRT2A / FRT2A, Ubi-RFP females, NosFlp;; *iml1^ko^,* FRT2A / FRT2A, Ubi-RFP females, were used to generate *mio^ko2^, ragA*^11^ *, ragc*^13^ *, nprl3^1^, iml1*^13^ mutant clones, homozygous clones were marked by absence of single GFP or RFP expression. Ovaries were dissected in Schneider’s medium from 2 days females after fed or starvation treatment next used for staining or live imaging assays.

### *Drosophila* germ cell Mass Spectrometric analyses and Co-Immunoprecipitation

Iml1-1xV5 and Nprl2-5xT7 knockin female flies were treated with regular fly food with yeast and/or starved for 12 hours, 30 pairs of ovaries were dissected on ice, homogenized in by TLB buffer (25 mM Tris-HCl pH 7.5, 5 mM magnesium chloride, 0.5% Triton. 1x protease inhibitors) ^42^, containing Complete Protease Inhibitors (Roche), Millipore T7 Tag Affinity Purification Kit and MBL v5 Affinity Purification kit were used for pulling down T7 and v5 tagged protein according to manufacturer’s instruction, the beads were sent to Taplin Biological Mass Spectrometry Facility and analyzed by mass spectrometry.

Iml1-1xV5, Nprl2-5xT7 knockin and RagC-GFP genetic insertion wild type and *wdr24* mutant female flies were treated with regular fly food with yeast and/or starved for 12 hours, 30 pairs of ovaries were dissected on ice, homogenized in TLB buffer. The samples were subjected to western blot analysis.

### *Drosophila* germ cell lysosome Immunoisolation

*Drosophila* germ cell lysosome immunoisolation was carried out using magnetic beads as previously described (^55^). Briefly, 30 pairs of Lamp1-3mcherry genomic insertion Drosophila ovaries in WT and *wdr24* mutant were homogenized in KPBS (136 mM KCl, 10 mM KH2PO4, adjusted to pH 7.25 with KOH). Fractions were incubated with anti-RFP magnetic beads with rotation at 4°C for 4 h. Beads were washed three times with KPBS. Bound lysosomes were subjected to SDS-PAGE and immunoblotting.

### Lysosomal FRAP Assay in Drosophila Germline

The FRAP experiments were performed on stage 6 egg chambers in Drosophila ovaries as described previously (Shu dev cell). Briefly, flies were cultured in standard media for 2-3 days before dissection. The ovaries were dissected in Schneider’s medium supplemented with 10% FBS, 10 μg/ml insulin, 5 μM nocodazole and 1μg/ml Hoechst. For starvation, the fed flies were transferred to 1% agarose/H2O media for 16hrs. The ovaries were dissected in amino acid starvation media as previously described (Wilson et al., 2004) and supplied with 5 μM nocodazole and 1mg/ml Hoechst 33342. 100% laser was used for lysosomal photobleaching, images were acquired at 10 second intervals for 4 mins. Mobile fraction was measured by using the last 7 frames to estimate the plateau. Mobile fraction (MF) ≈ (I (plateau) - I0) / (I (pre-bleach) - I0). I0: The fluorescence intensity immediately after photobleaching.

### Lysosomal FRAP Assay in HeLa cell

FRAP experiments were performed on HeLa cells as described previously (Shu et al, 2020). The recovery fraction was calculated by fitting the FRAP fluorescence recovery curve into the model Y=Y0 + (Plateau-Y0)*(1-exp(-K*x)) through using one-phase association

## Quantification and statistical analysis

The details show in figure legends. All represent data from at least three independent experiments. Statistical comparisons were made using Unpaired Student’s t-test provided by GraphPad Prism 10 software.

## Key resources table

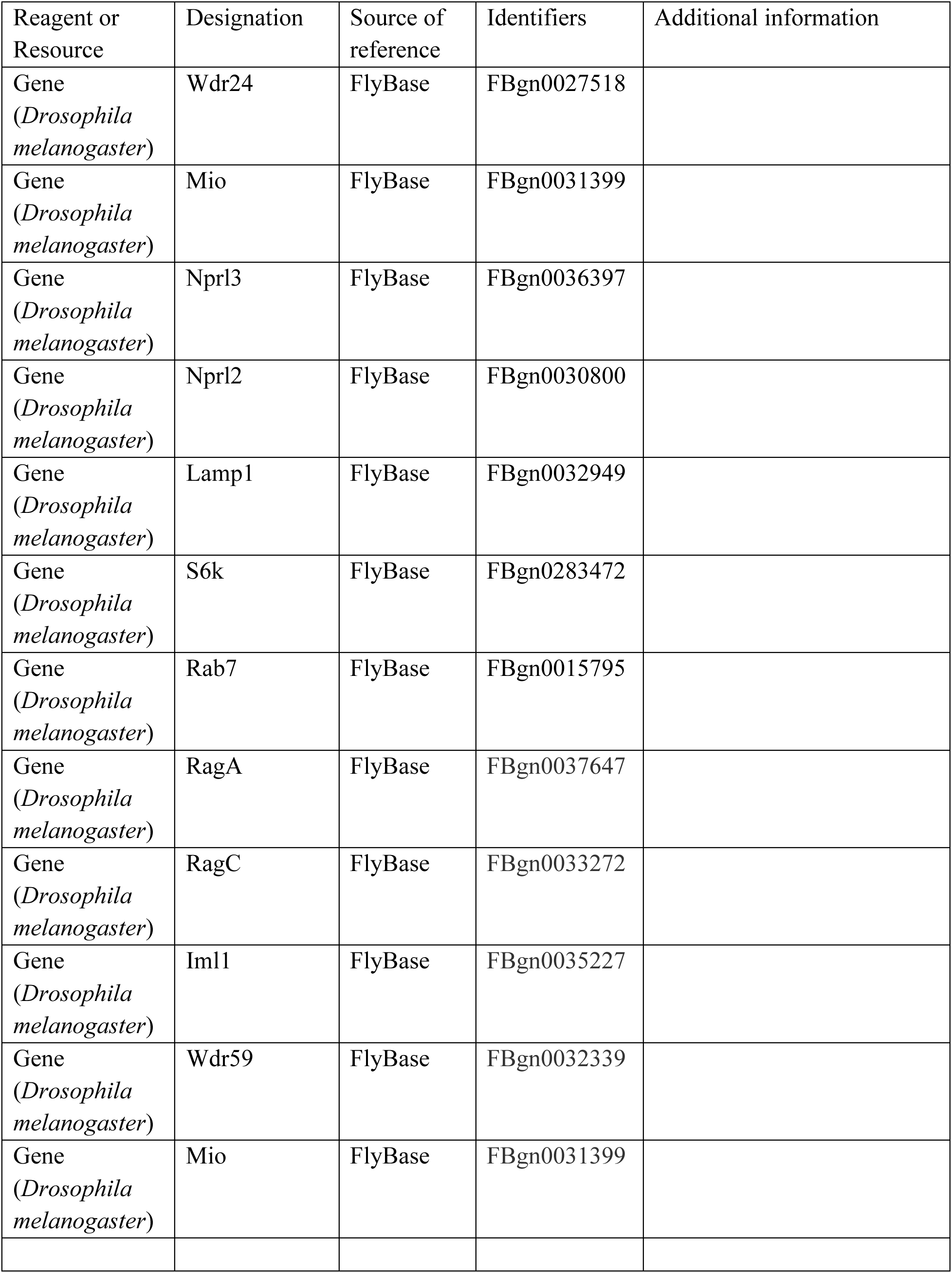
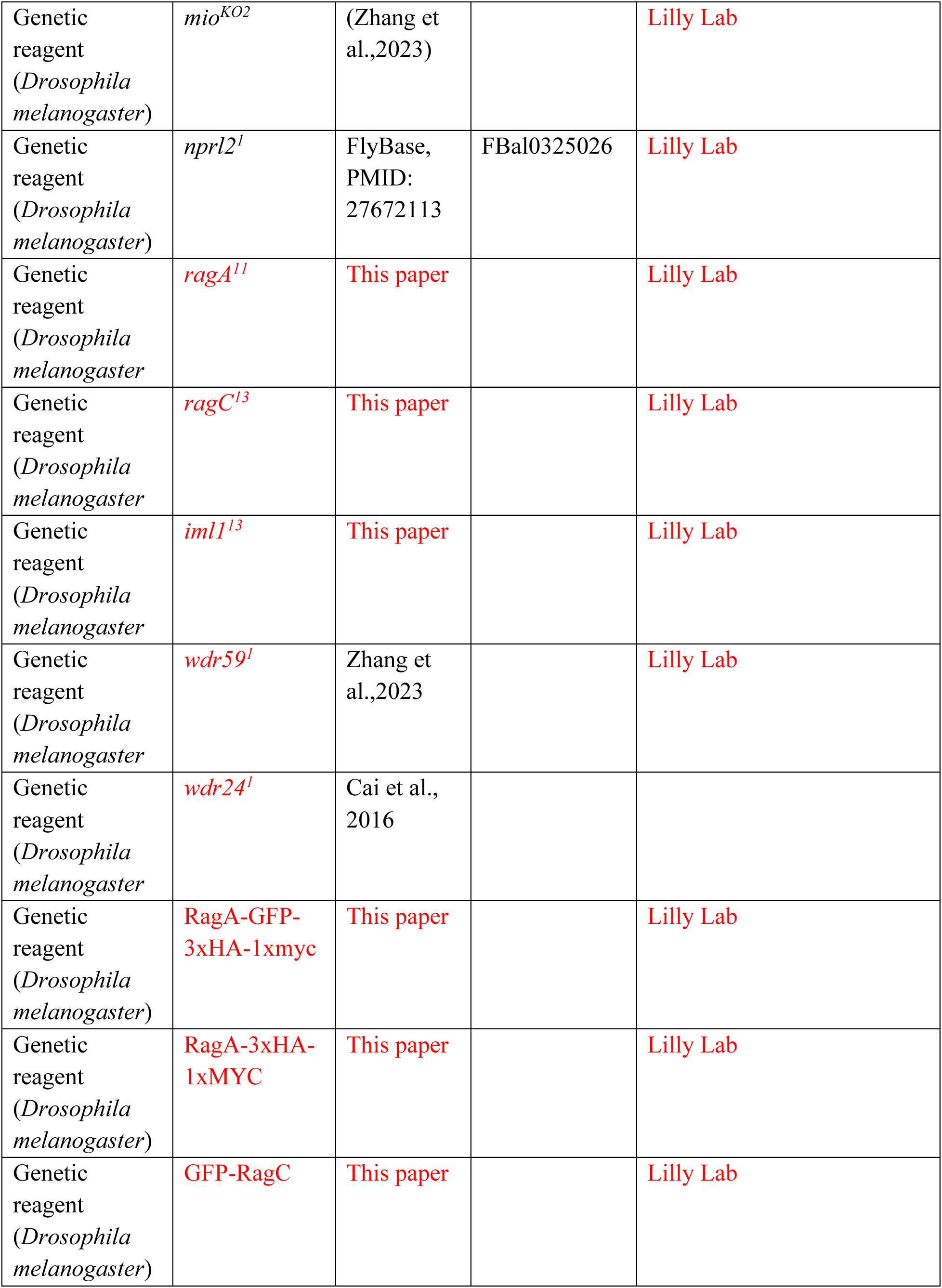
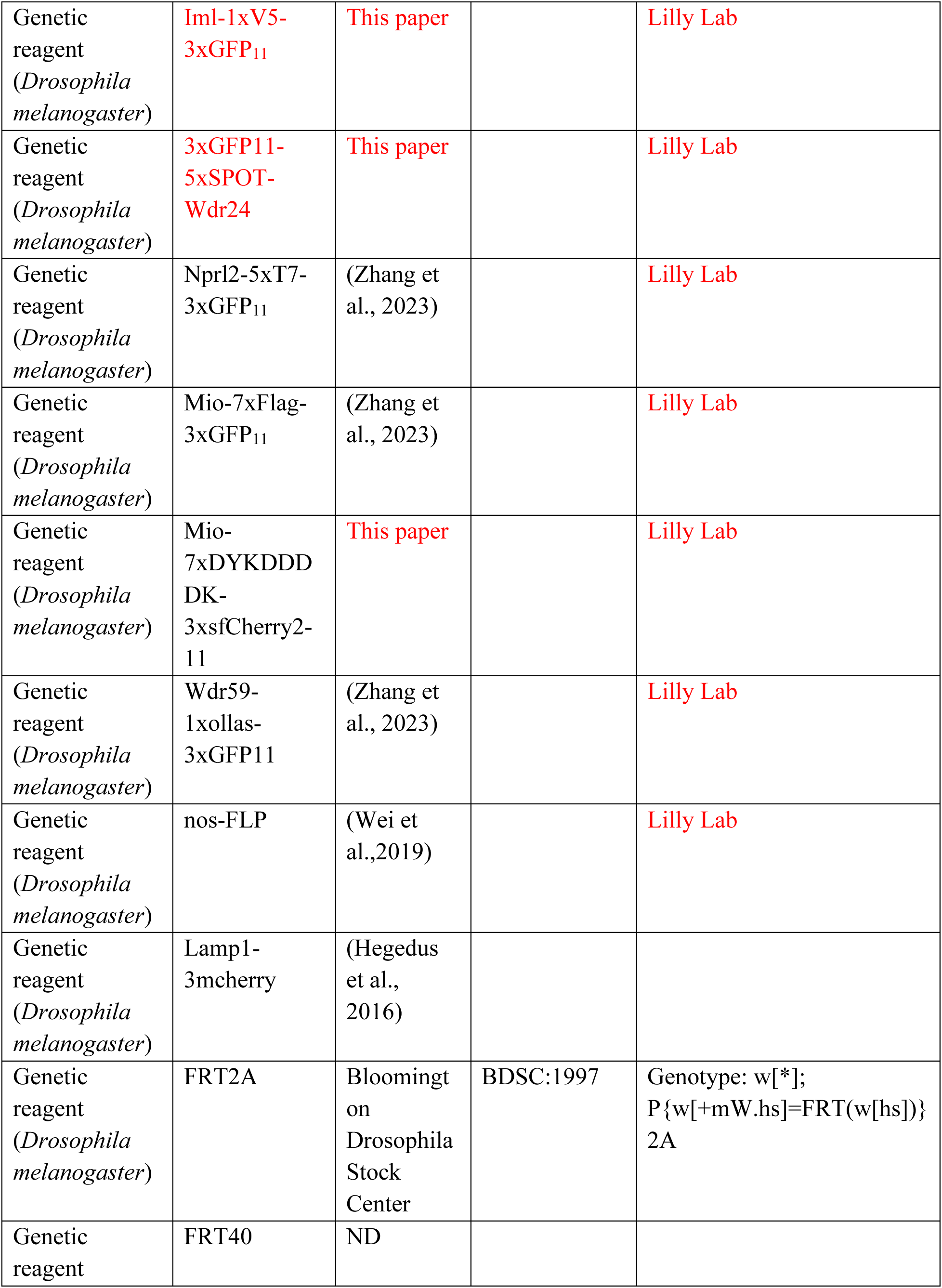
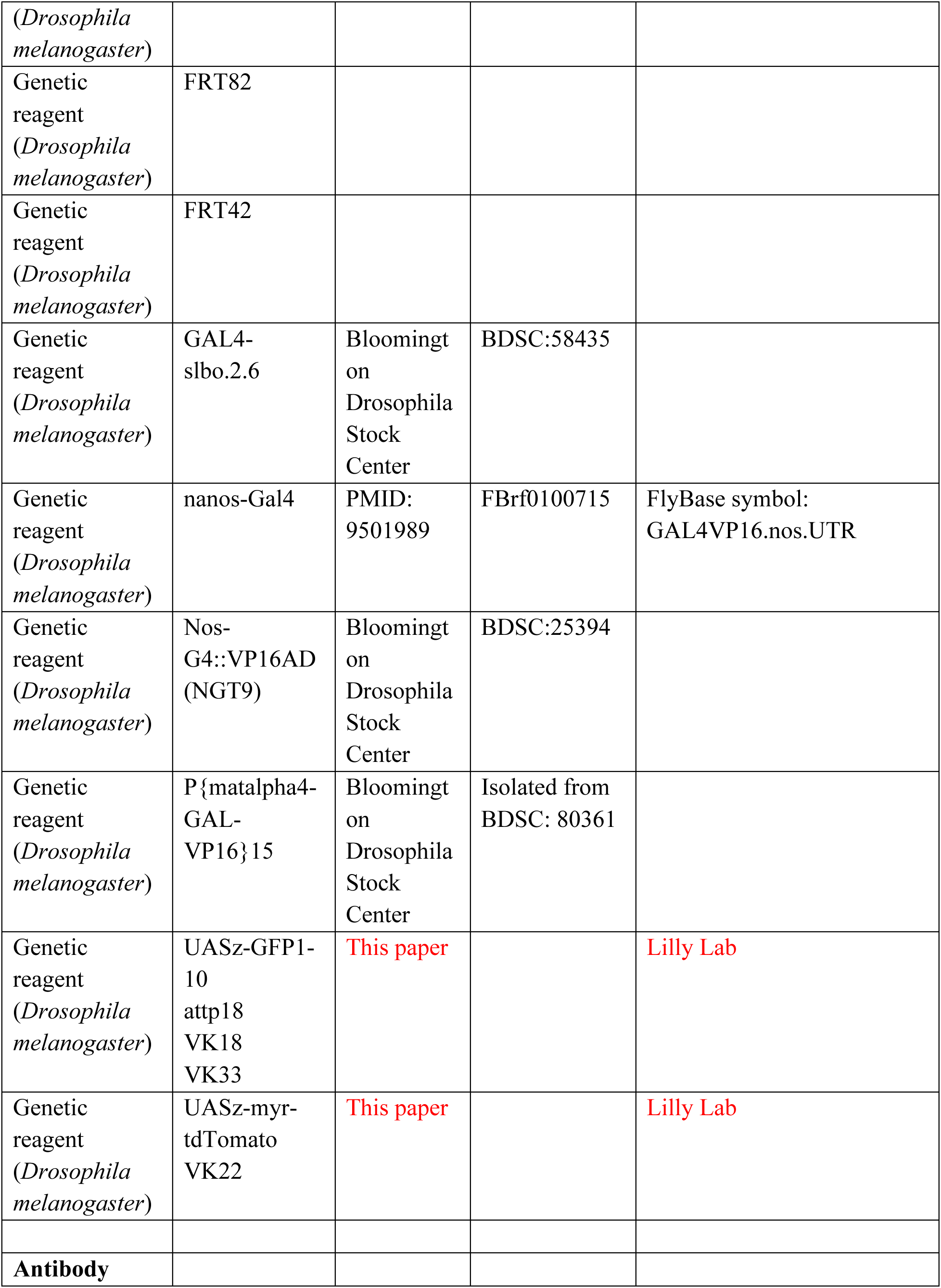
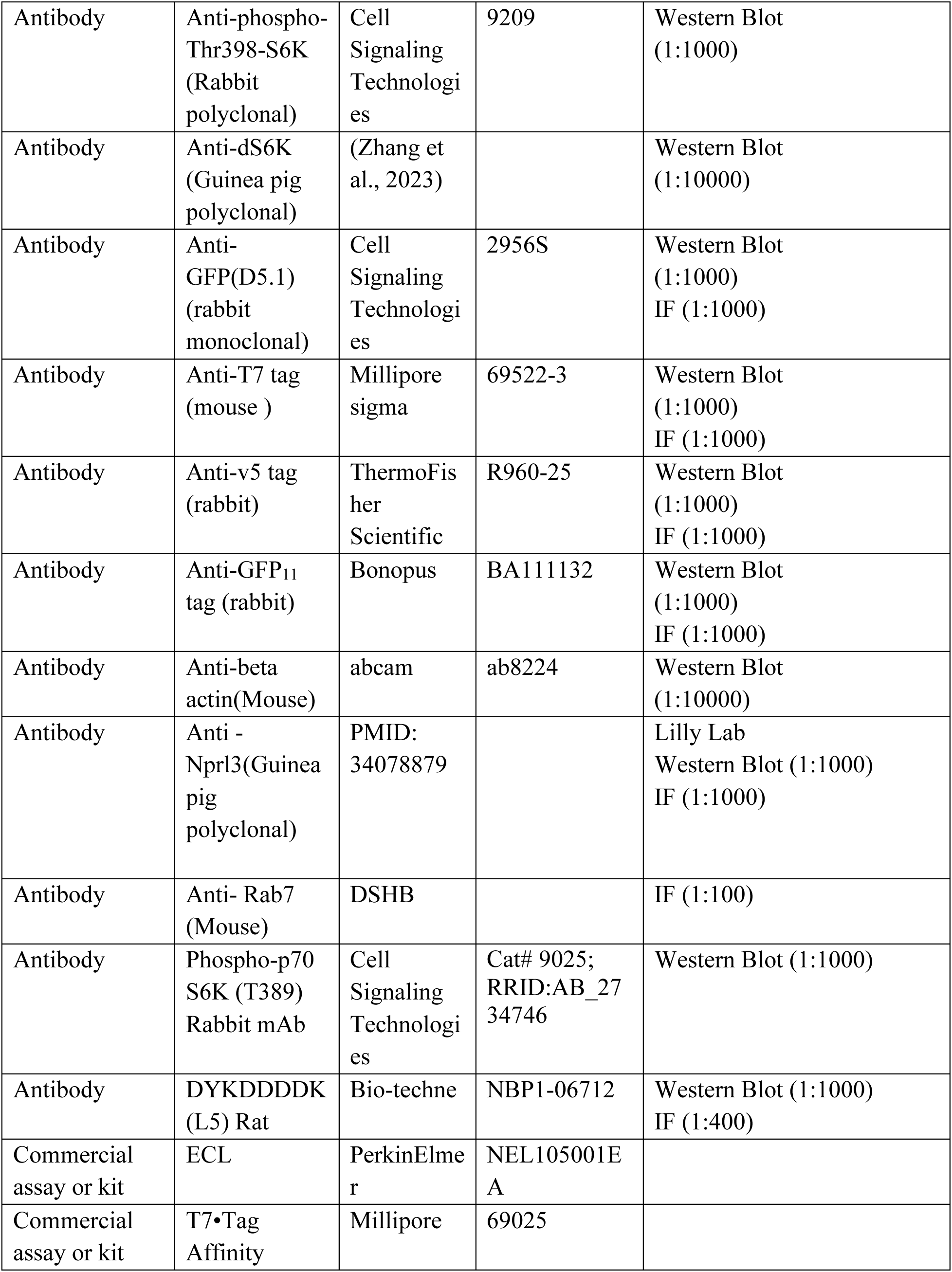
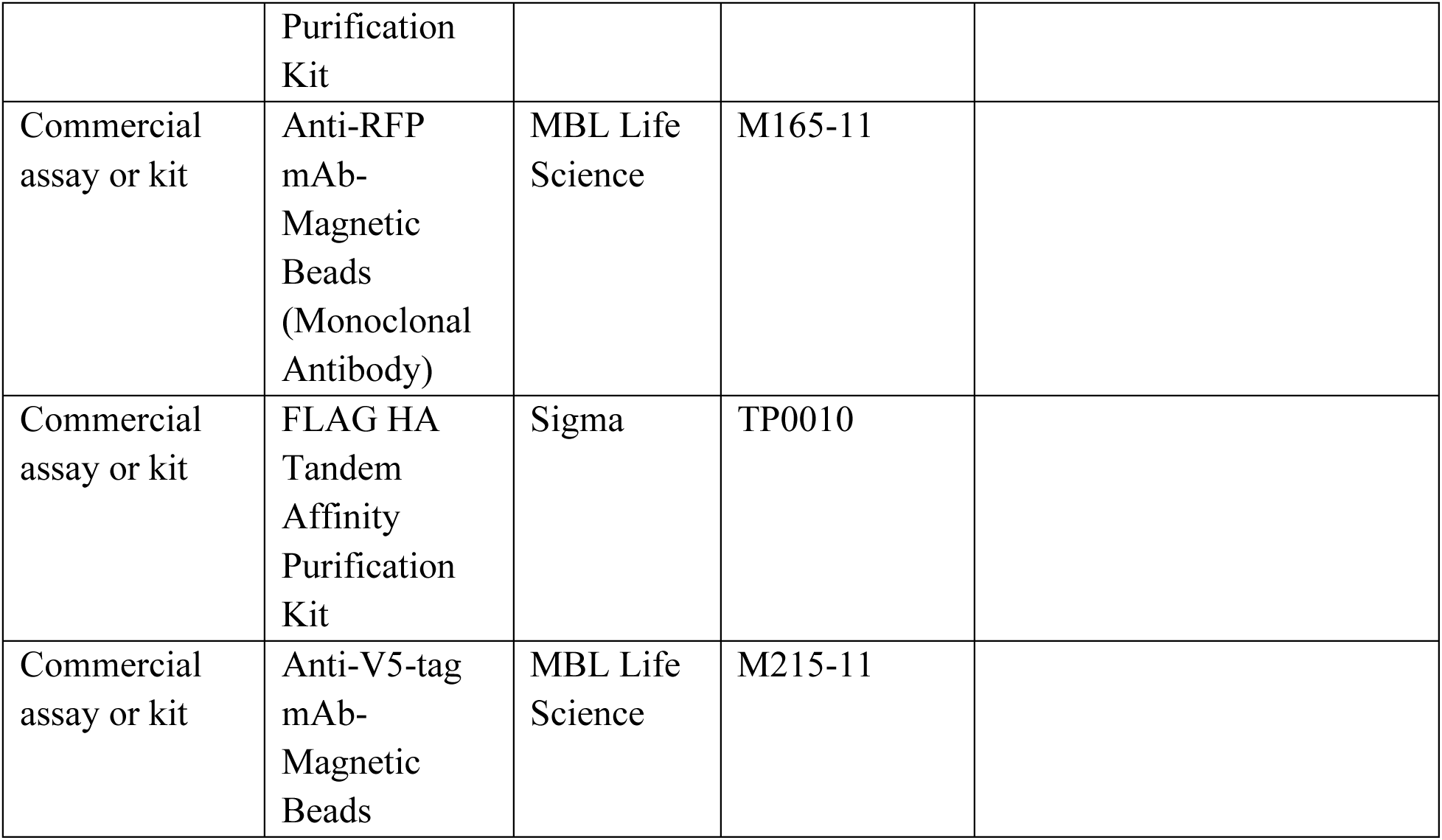

---

**Table.**
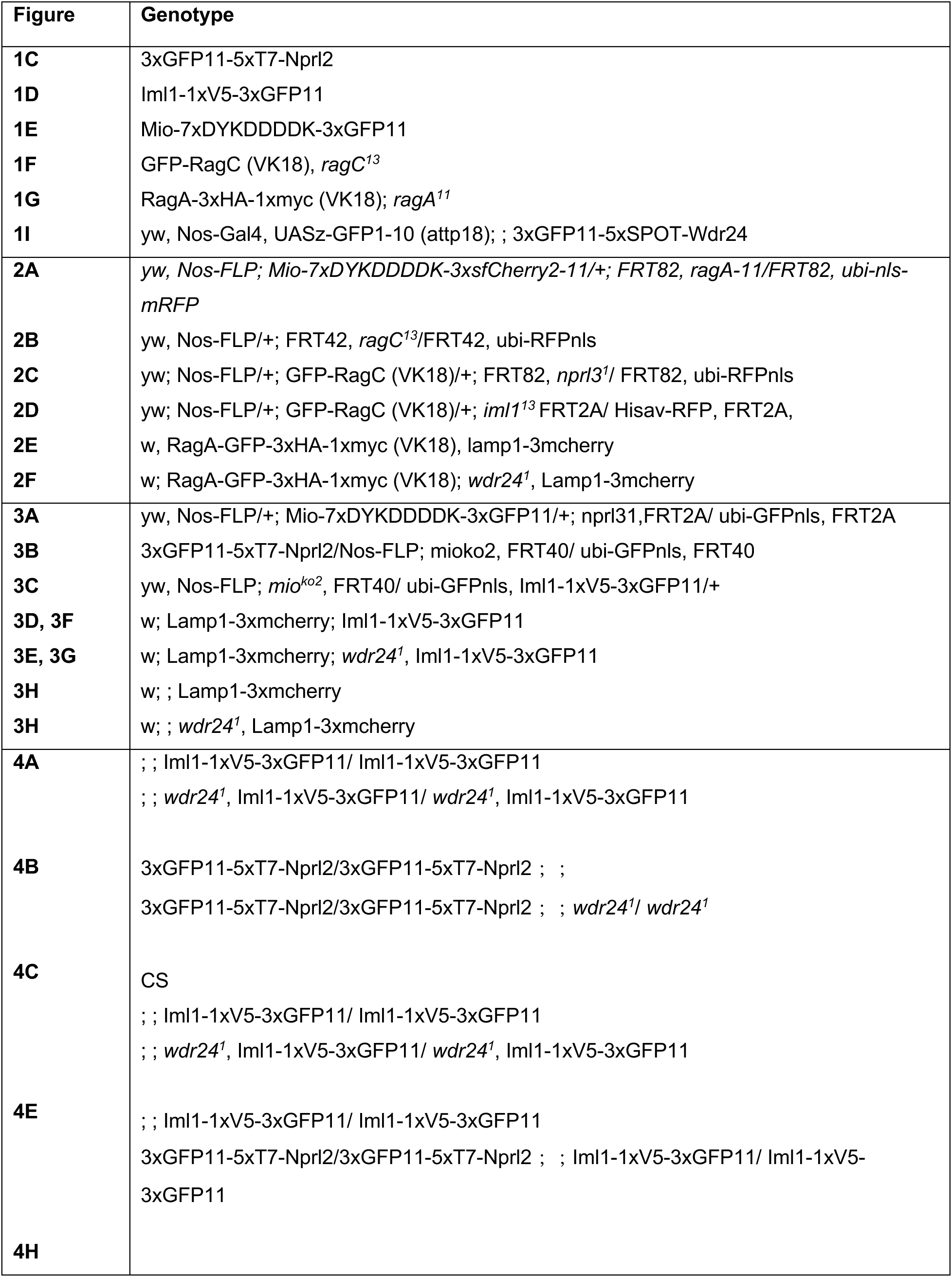
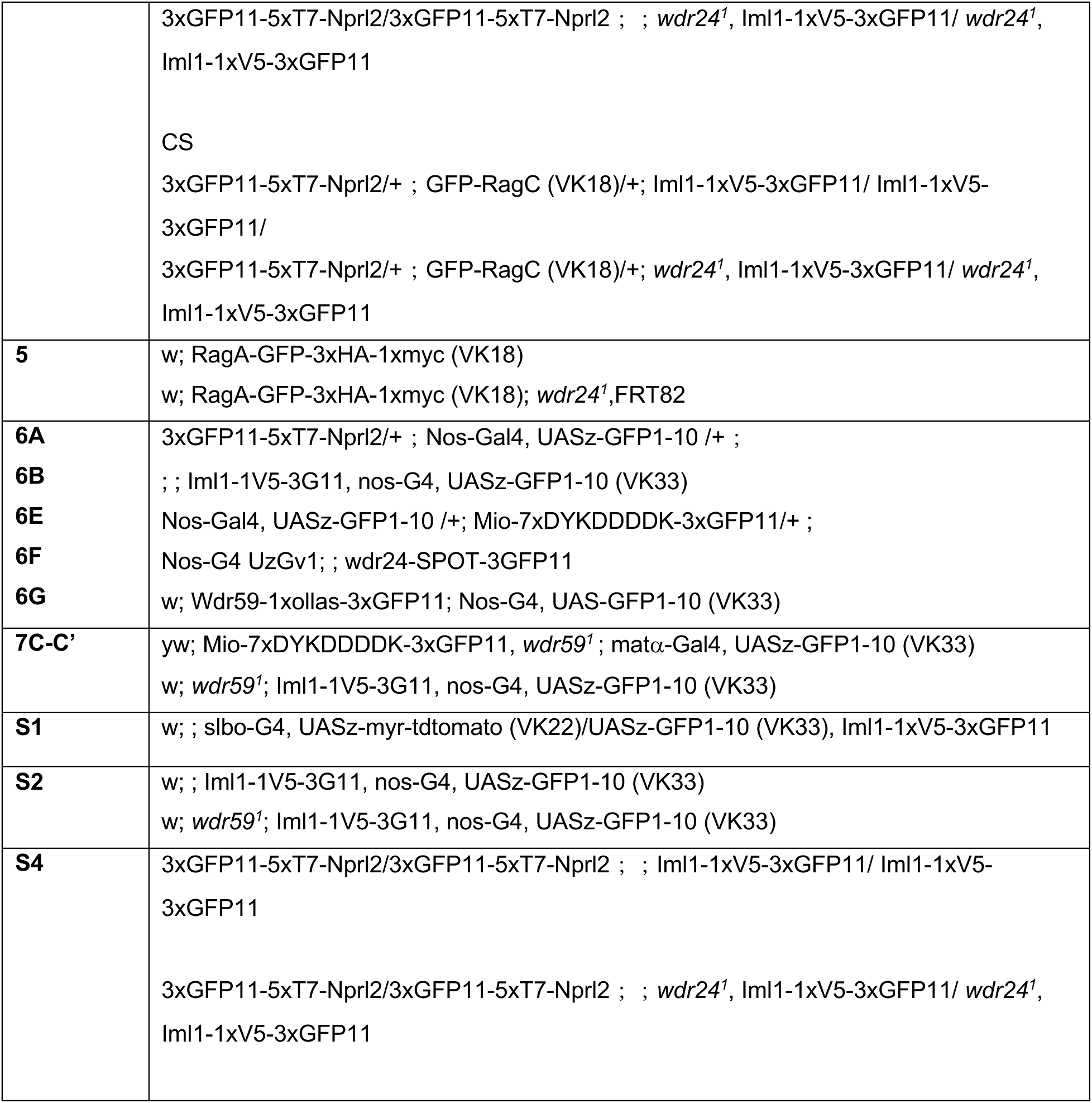
Full Genotypes of Flies Shown in Main Figures and Supplemental Figures.

