## Supplemental Figures for "GATOR2 regulates the nutrient-dependent recruitment of GATOR1 to lysosomes"

#### Figure S1

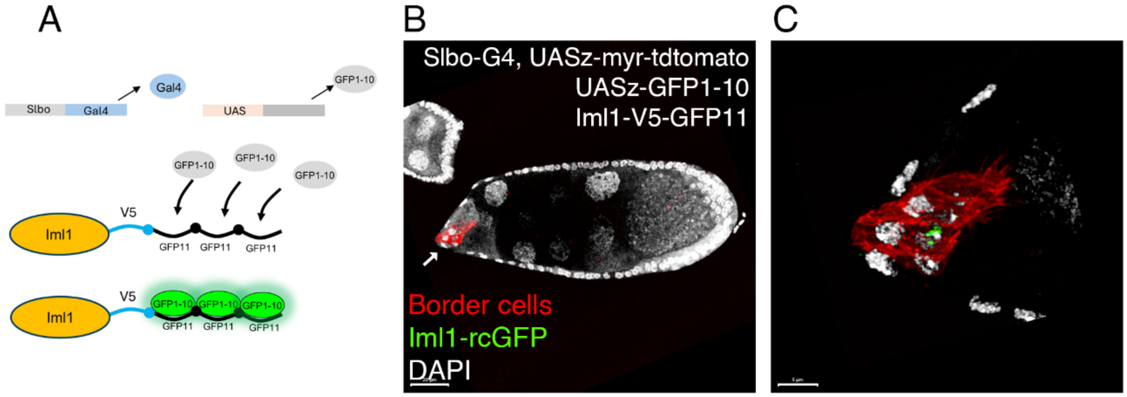


**Supplemental Figure 1. Iml1-rcGFP in border cell lysosomes in stage 9 egg chamber.**

(A) Schematic representation of Iml1-specific labeling in border cells using the split-GFP system.

(B) Membrane-tethered tdTomato (red, arrow) and GFP1–10 were co-expressed in border cells using the border cell–specific driver slbo-Gal4. Knock-in Iml1 (green) was reconstituted via GFP complementation. DNA was labeled with DAPI (white).

(C) Reconstituted Iml1–GFP signal is specifically detected in border cells, demonstrating tissue specific labeling capability. Scale bar for (B) 20 μm. (C) 5μm.

### Figure S2

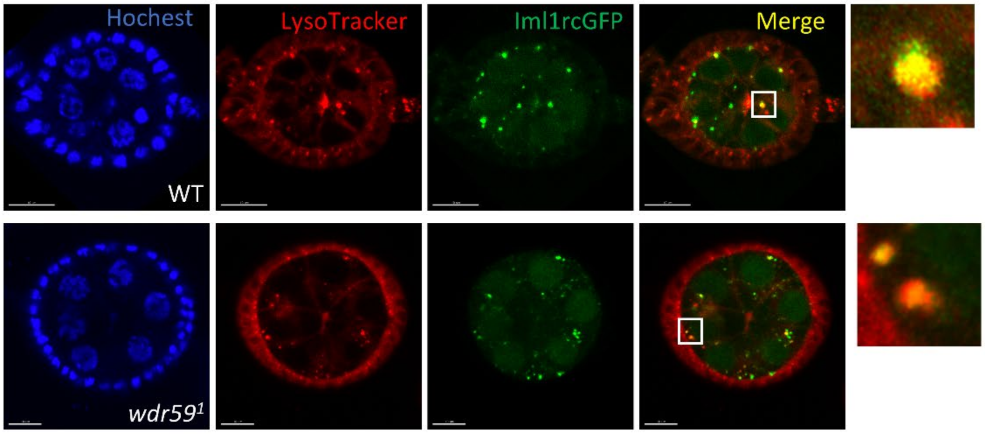


**Supplemental Figure 2. Wdr59 is not required for the localization of GATOR1 to lysosomes or autolysosomes.**

Live cell imaging of Drosophila egg chambers from WT and *wdr59¹* mutant females cultured on standard fly medium. Iml1-rcGFP colocalizes with the lysosomal marker LysoTracker. Scale bar: 10 μm.

### Figure S3

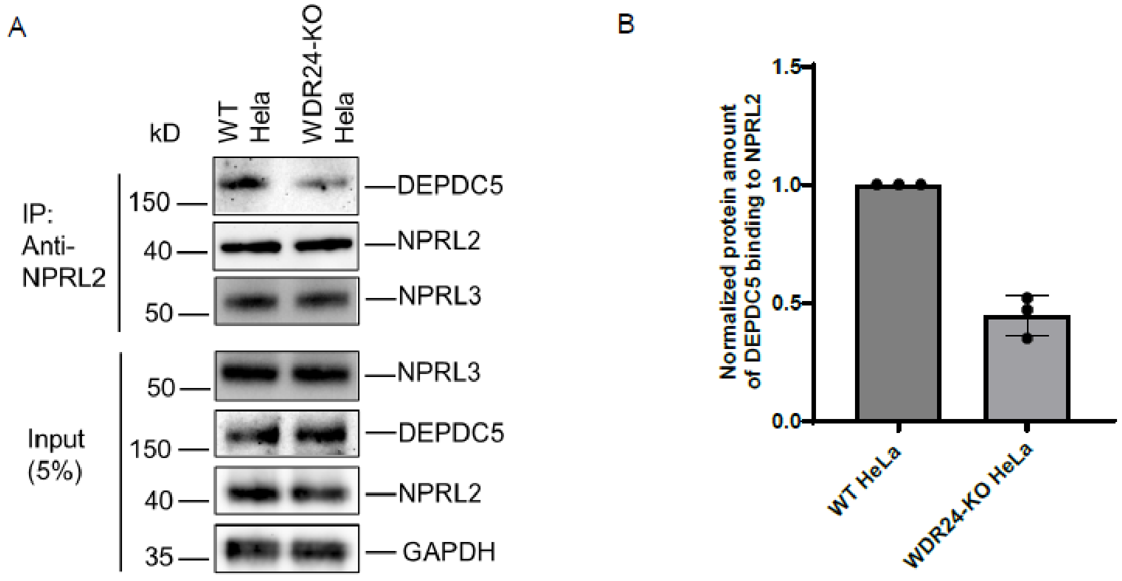


**Supplemental Figure 3. The association between NPRL2 and DEPDC5 decreased in WDR24-KO HeLa cells.**

(A). Endogenous co-immunoprecipitation using anti-NPRL2 antibody to pull down NPRL2, showed that the amount of DEPDC5 binding to NPRL2 decreased in WDR24-KO HeLa cells, compared to WT HeLa cells.

(B). Quantification of the protein amount of DEPDC5 binding to NPRL2 from three biological replicates. Protein amounts of DEPDC5 in WT and WDR24-KO cells from the “IP” group were first normalized to the DEPDC5 of each cell in the “Input” group, respectively; then the amount from the WDR24-KO cells were normalized to the WT HeLa cells group.

### Figure S4

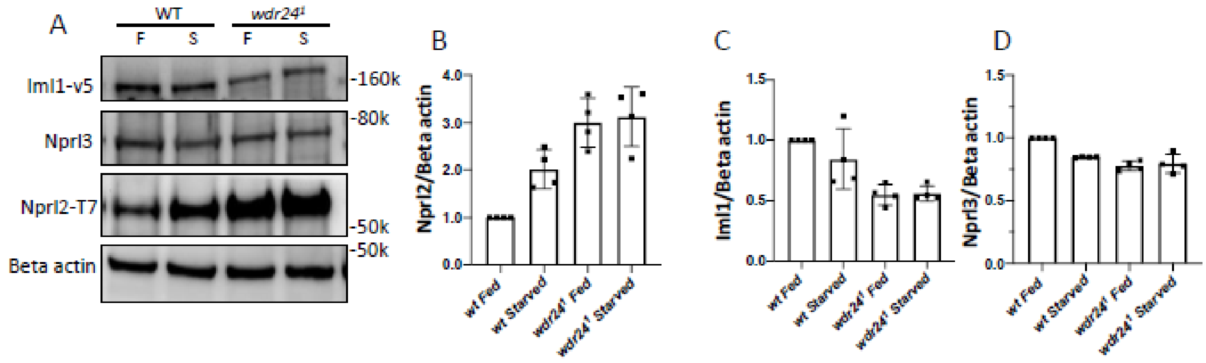


**Supplemental Figure 4. GATOR2 decreases the stability of Nprl2.**
(A-D) Western blot of GATOR1 components, Iml1, Nprl3 and Nprl2 total protein levels from ovarian lysates prepared from fed and starved(12h) WT and *wdr24*$^{1}$ females. (B-D) Quantification of Nprl2, Iml1 and Nprl3 levels relative to Beta actin from(A).
